# Computational modeling of drug response identifies mutant-specific constraints for dosing panRAF and MEK inhibitors in melanoma

**DOI:** 10.1101/2024.08.02.606432

**Authors:** Andrew Goetz, Frances Shanahan, Logan Brooks, Eva Lin, Rana Mroue, Darlene Dela Cruz, Thomas Hunsaker, Bartosz Czech, Purushottam Dixit, Udi Segal, Scott Martin, Scott A. Foster, Luca Gerosa

## Abstract

Purpose: This study explores the potential of preclinical *in vitro* cell line response data and computational modeling in identifying optimal dosage requirements of pan-RAF (Belvarafenib) and MEK (Cobimetinib) inhibitors in melanoma treatment. Our research is motivated by the critical role of drug combinations in enhancing anti-cancer responses and the need to close the knowledge gap around selecting effective dosing strategies to maximize their potential. Results: In a drug combination screen of 43 melanoma cell lines, we identified unique dosage landscapes of panRAF and MEK inhibitors for NRAS vs BRAF mutant melanomas. Both experienced benefits, but with a notably more synergistic and narrow dosage range for NRAS mutant melanoma. Computational modeling and molecular experiments attributed the difference to a mechanism of adaptive resistance by negative feedback. We validated *in vivo* translatability of *in vitro* dose-response maps by accurately predicting tumor growth in xenografts. Then, we analyzed pharmacokinetic and tumor growth data from Phase 1 clinical trials of Belvarafenib with Cobimetinib to show that the synergy requirement imposes stricter precision dose constraints in NRAS mutant melanoma patients. Conclusion: Leveraging pre-clinical data and computational modeling, our approach proposes dosage strategies that can optimize synergy in drug combinations, while also bringing forth the real-world challenges of staying within a precise dose range.

**Simple Summary:** Combining drugs is crucial for enhancing anti-cancer responses. However, the potential of pre-clinical data in identifying suitable combinations and dosage is often underutilized. In this study, we leverage preclinical *in vitro* cell line drug response data and computational modeling of signal transduction and of pharmacokinetics to elucidate distinct dose requirements for the combination of pan-RAF and MEK inhibitors in melanoma. Our findings reveal a more synergistic, but narrower dosing landscape in NRAS vs BRAF mutant melanoma, which we linked to a mechanism of adaptive resistance through negative feedback. Further, our analysis suggests the importance of drug dosing strategies to optimize synergy based on mutational context, yet highlights the real-world challenges of maintaining a narrow dose range. This approach establishes a framework for translational investigation of drug responses in the refinement of combination therapy, balancing the potential for synergy and practical feasibility in cancer treatment planning.

## 1. Introduction

Cancer is a disease marked by abnormal cell growth and the potential to spread and cause death. Despite its complexities, cancers often carry vulnerabilities that make them susceptible to targeted treatments [1–3]. Precision medicine provides a promising approach to exploit these vulnerabilities and effectively kill cancer cells. However, designing effective targeted therapies is not straightforward. The dynamic nature of cancer cells enables them to adapt and develop resistance mechanisms, often rendering single-drug treatments less effective [4,5]. As a response, the medical field has turned towards combined drug regimens, simultaneously targeting multiple vulnerabilities in cancer cells. Identifying effective drug combinations, however, is only one part of the puzzle. The dosing regimes of these combinations that yield maximal benefit while maintaining tolerability must also be determined. Current approaches to delineate these aspects often fall short.

I*n vitro* drug screens using cancer cell lines represent a primary tool for identifying drug combinations that act beneficially on lines exhibiting traits of interest [6,7]. Typically, changes in cell viability are measured in response to the serial dilution of two drugs, also called a drug dose-response matrix, and the benefits of combining drugs is quantified based on principles such as Highest Single Agent (HSA), Bliss independence, Loewe additivity, and others [8,9]. These enable the computation of combination scores, which are used to rank the effectiveness of drug combinations with respect to single agents. A significant limitation in the use of combination scores is the inadequate consideration of the specific point in the dose-response landscape where benefits are observed, leading to the omission of drug dose from the benefit assessment. This can lead to an inaccurate assessment of clinical potentials and a mischaracterization of biomarkers, particularly in situations where cancer populations exhibit responses at distinct effective dose ranges.

The reasons for these limitations are both practical and conceptual. A practical limitation is the lack of computational frameworks for easily manipulating large-scale dose-response data and extracting dose-specific information. While tools that adhere to FAIR software principles have been recently developed [10,11], they still lack mature capabilities for extracting and analyzing response data at the (free) drug concentrations determined by pharmacokinetics in the clinic [12]. A more profound conceptual limitation is the unclear translatability of *in vitro* drug responses to *in vivo* settings. The primary strategies used are either qualitative, such as benchmarking exposures to single-point i*n vitro* metrics like the half-maximal inhibitory concentration (IC50) values, or require extensive datasets and efforts, as in mechanistic modeling [13] or machine learning [14]. Recently, success has been reported in using *in vitro* growth rate inhibition values with pharmacokinetic parameters to estimate *in vivo* drug response [15,16], but these results were limited to single-agent response. Improving the frameworks for drug dose-response analysis and testing the translatability of *in vitro* drug combinations to *in vivo* is required to exploit the full potential of pre-clinical data.

While dose-response experiments with cell lines provide insightful data on drug impact, their phenomenological nature limits mechanistic understanding. Thus, methods able to link dose-response data to molecular measurements and information on protein structures and networks are needed. Increasingly, computational dynamic models — mathematical representations of molecular networks — are being deployed to elucidate these mechanisms [4,17]. Due to its role in cancer and advanced molecular understanding, the MAPK signaling pathway has been the focus of current developments of computational models of drug response [18–22]. These models are perpetually updated to incorporate new conditions and advancements in the understanding of oncogenic signaling. A necessary development is the use of these models to explain variations seen in drug responses based on traits of interest, such as mutational status, and link phenotypes to mechanistic insights at the clinically relevant dose. The promise is that these models can generalize correlative trends based on theoretical reasonings and provide molecular insights that can be experimentally verified.

In this study, we deploy a framework that combines preclinical *in vitro* cell line drug response data and computational modeling of signal transduction and pharmacokinetics to unravel the dose requirements of pan-RAF and MEK inhibition for melanoma treatment. The development of RAF inhibitors has seen many advancements with initial, first-generation inhibitors showing effectiveness against active RAF monomers such as BRAF V600E [23]. The primary limitation of these inhibitors is their inability to block, and sometimes even paradoxically enhance, RAF dimer signaling. As a result, the inhibitors are ineffective against prevalent mutations like NRAS Q61, which signal through RAF dimers, and are liable to escape mechanisms through RAF dimer signaling [24]. This has spurred the development of several small-molecule ATP-competitive panRAF inhibitors, such as Belvarafenib [25], which are capable of targeting RAF dimers and are currently in clinical trials. Bolstered by robust preclinical evidence [26–28], in the clinic panRAF inhibitors are being combined with MEK inhibitors to achieve stronger pathway suppression and avoid mechanisms of resistance [trials: NCT03284502, NCT04417621, NCT03905148, NCT04249843 and NCT03429803]. However, the ways in which these drugs inhibit activity under the two major activating mutations in melanoma, BRAF V600E and NRAS Q61 hotspot mutations, and the corresponding drug dose landscape are still being explored. To this end, we apply our approach in the hopes of unraveling how this drug combination impacts different mutational contexts and identifying effective drug regimens for clinical use.

## 2. Materials and Methods

### 2.1 Drug combination screen

Screening Drugs Management and Quality Control. Drugs were obtained from in-house synthesis or purchased from commercial vendors. A fully automated transfer system by Nova Technology (Innovate Engineering) was used to transfer material from a dry library, solubilize with DMSO, and log the solutions into our compound management system. A high throughput liquid chromatography mass spectrometry/ultraviolet absorbance/charged aerosol detector/chemiluminescent nitrogen detector (LCMS/UV/CAD/CLND) system was used to verify the identity, purity, and concentration of drugs used in the gCSI screens. The LCMS/UV/CAD/CLND system consisted of an LCMS/UV system (Shimadzu) with LC-30AD solvent pump, 2020 MS, Sil-30AC autosampler, SP-M30A UV detector, and CTO-20A column oven; a Corona Veo RS CAD (Thermo Scientific); and a model 8060 CLND. Drugs with lower than 80% purity and 20% below expected concentration were excluded. An Echo 555 acoustic drop ejection (ADE) liquid handler (Labcyte) was fully integrated in the ultra-high-throughput screening uHTS system to dispense DMSO solubilized compounds (Dawes et al., 2016). Nine-point dose–response curves at 1:3 dilution were generated using ADE as a means of transferring library compounds at ultra-low volume (in nanolitre scale) to achieve direct dilution of compounds. Starting doses for Vemurafenib, Belvarafenib and Cobimetinib were 10, 10 and 5 µM, respectively. The uHTS system delivered assay-ready daughter plates at 31,000 concentration. A DMSO backfill step was performed to achieve an equal volume of DMSO in each well. Assay-ready drug plates were stored at -80 C until the day of compound addition and subjected to a single freeze-thaw cycle. The use of ADE technology limited the final DMSO concentration in assay plates to 0.1%, which was shown to have a negligible effect on cell growth. Seeding densities were optimized for each cell line to obtain 70-80% confluence after 6 days. The cells were plated into 384 well plates (Greiner, 781091) and then treated with compound the following day in a final DMSO concentration of 0.1%. The relative numbers of viable cells were measured by luminescence using CellTiter-Glo (Promega, G7573).

### 2.2 Higher drug-dose resolution combination responses

We generate higher drug-dose resolved 10×10 drug combination responses centered around clinically relevant doses for 5 cell lines: A375, IPC-298, MEL-JUSO, SK-MEL-2 and SK-MEL-30. Seeding densities were optimized to obtain 70-80% confluence after 6 days. Cells were seeded into 384-well plates 24 hours prior to compound addition, and treated with compound the following day (final DMSO concentration 0.1%). Compound stocks, 10 mM in DMSO, supplied by Genentech Compound Management. Belvarafenib and Cobimetinib were dosed using an HP 300 automatic dose dispenser as a 10 x 10 combinatorial drug matrix with serial dose dilutions starting from 1 to 0.002 µM for Belvarafenib and 0.5 to 0.002 µM for Cobimetinib. After 120 hours, relative numbers of viable cells were measured using Cell Titer-Glo (Promega, G7573).

### 2.3 Western Blots

Anti-MEK1 (12671, WB 1:1,000), anti-pMEK (S217/S221) rabbit mAb (41G9) (9154, WB 1:1,000), anti-ERK (9107, WB 1:1,000), anti-pERK (T202/Y204) (9101, WB 1:1,000), purchased from Cell Signaling Technology. IR-conjugated secondary antibodies, Goat anti-Mouse 680LT (926-68020, WB: 1:10,000) and Goat anti-Rabbit 800CW (926-32211, WB: 1:10,000) purchased from Li-Cor. All westerns were scanned on Li-Cor Odyssey CLX using duplexed IR-conjugated secondary antibodies.

SK-MEL-28, A-375, and SK-MEL-2 were obtained from ATCC. IPC-298 and MEL-JUSO were obtained from DSMZ. Cell lines were maintained in the recommended media and supplemented with 10% heat-inactivated FBS (HyClone, SH3007003HI), 1X GlutaMAX (Gibco, 35050-061), and 1X Pen Strep (Gibco, 15140-122).

Immunoblotting was performed using standard methods. Cells were briefly washed in ice-cold PBS and lysed in the following lysis buffer (1% NP40, 50 mM Tris, pH 7.8, 150 mM NaCl, 5 mM EDTA) plus protease inhibitor mixture (Complete mini tablets; Roche Applied Science, 11836170001) and phosphatase inhibitor mix (ThermoFisher Scientific, 78420). Lysates were centrifuged at 15,000 rpm for 10 minutes at 4 °C and the protein concentration was determined by BCA (ThermoFisher Scientific, 23227). Equal amounts of protein were resolved by SDS-PAGE on NuPAGE, 4-12% Bis-Tris Gels (ThermoFisher Scientific, WG-1403) and transferred to nitrocellulose membrane (Bio-Rad, 170-4159). After blocking in blocking buffer (Li-Cor, 927-40000), membranes were incubated with the indicated primary antibodies and analyzed by the addition of secondary antibodies IRDye 680LT Goat anti-Mouse IgG (Li-Cor, 926-68050) or IRDye 800CW Goat anti-Rabbit IgG (Li-Cor, 926-32211). The membranes were visualized on a Li-Cor Odyssey CLx Scanner.

### 2.4 Immunofluorescence and high-content imaging

Cells were washed twice with 1x PBS and fixed with 4% paraformaldehyde (PFA) for 15 min at 25 °C. To remove PFA, cells were washed with 1x PBS three times, and PFA was quenched by incubating cells with 50 mM NH_4_Cl for 10 min at 25 °C. Cells were then rinsed twice with PBS and permeabilized with ice-cold methanol for 10 min at -20 °C. Following permeabilization, cells were first incubated with a blocking buffer for 1 hour at room temperature (1x PBS/ 5% normal serum/0.3% TritonX-100) followed by overnight incubation with the primary antibody against phospho-ERK (Cell Signaling Technology, catalog no. 4370S) at 1:800 dilution at 4 °C. The next day, cells were washed three times with 1x PBS and incubated for one hour at room temperature with the secondary antibody (Jackson ImmunoResearch Laboratories, catalog no. 711-606-152). To stain the nucleus and cell body, cells were incubated with NucBlue™ Fixed Cell ReadyProbes™ Reagent (Catalog number: R37606) and HCS CellMask™ Blue Stain (Catalog number: H32720) for 20 min at room temperature. Finally, cells were washed three times with 1X PBS, and imaged on the Opera Phenix HCS machine (PerkinElmer) using the 40X water immersion objective using confocal modality. Analysis and quantification were conducted on Harmony (PerkinElmer) software.

### 2.5 Tumor volume experiments in xenografts

G03083045.23-6 (free base of GDC-5573, Lot 23-6; hereafter referred to as Belvarafenib) was provided to Genentech as a solution at concentrations of 3.3 mg/mL and 6.6 mg/mL (expressed as free-base equivalents) in 5% dimethyl sulfide/5% Cremophor EL. Cobimetinib (GDC-0973, Lot 150-10) was provided by Genentech as a solution at concentrations of 1.1 mg/mL (expressed as free-base equivalents) in 0.5% (w/v) methylcellulose/0.2% Tween 80™. All concentrations were calculated based on a mean body weight of 22 g for the NCR.nude mouse strain used in this study. The vehicle controls were 5% dimethyl sulfide/5% Cremophor EL and 0.5% (w/v) methylcellulose/0.2% Tween 80™. Test articles were stored in a refrigerator set to maintain a temperature range of 4-7 °C. All treatments and vehicle control dosing solutions were prepared once a week for three weeks.

Female NCR.nude mice that were 6-7 weeks old were obtained from Taconic Biosciences (New York) weighing an average of 22 g. The mice were housed at Genentech in standard rodent micro-isolator cages and were acclimated to study conditions at least 3 days before tumor cell implantation. Only animals that appeared to be healthy and that were free of obvious abnormalities were used for the study.

Human melanoma IPC-298 cells were obtained from the ATTC (Rockville, MD) harbor NRAS Q61L mutation. Cells were cultured in vitro, harvested in log-phase growth, and resuspended in Hank’s Balanced Salt Solution (HBSS) containing Matrigel (BD Biosciences; San Jose, CA) at a 1:1 ratio. The cells were then implanted subcutaneously in the right lateral thorax of 140 NCR.nude mice. Each mouse was injected with 20 * 10^6 cells in a volume of 100 mL. Tumors were monitored until they reached a mean tumor volume of 250-300 mm^3^. Mice were distributed into six groups based on tumor volumes with n=10 mice per group. The mean tumor volume across all six groups was 240 mm^3^ at the initiation of dosing.

Mice were given vehicles (100 µL 5% DMSO/5% CremEL and 100 µL 0.5% MCT), 15 mg/kg or 30 mg/kg Belvarafenib (expressed as free-base equivalents) and 5 mg/kg Cobimetinib (expressed as free-base equivalents). All treatments were administered on a daily basis (QD) orally (PO) by gavage for 21 days in a volume of 100 mL for Belvarafenib or Cobimetinib. Tumor sizes and mouse body weights were recorded twice weekly over the course of the study. Mice were promptly euthanized when tumor volume exceeded 2000 mm^3^ or if body weight loss was ≥ 20% of their starting weight.

All drug concentrations were calculated based on a mean body weight of 22 g for the NCR.nude mouse strain used in this study. The study design is summarized in Table S1. Tumor volumes were measured in two dimensions (length and width) using Ultra Cal-IV calipers (model 54 − 10 − 111; Fred V. Fowler Co.; Newton, MA) and analyzed using Excel, version 14.2.5 (Microsoft Corporation; Redmond, WA). The tumor volume was calculated with the following formula: Tumor size (mm^3^) = (longer measurement × shorter measurement^2) × 0.5. Animal body weights were measured using an Adventura Pro AV812 scale (Ohaus Corporation; Pine Brook, NJ). Percent weight change was calculated using the following formula: Body weight change (%) = [(current body weight/initial body weight) – 1) × 100] Percent animal weight was tracked for each individual animal while on study and the percent change in body weight for each group was calculated and plotted (**Figure S1**). A generalized additive mixed model (GAMM) was employed to analyze transformed tumor volumes over time. As tumors generally exhibit exponential growth, tumor volumes were subjected to natural log transformation before analysis. Changes in tumor volumes over time in each group are described by fits (i.e., regression splines with auto-generated spline bases) generated using customized functions in R version 3.4.2 (2017-09-28) (R Development Core Team 2008; R Foundation for Statistical Computing; Vienna, Austria).

For assessment of gene expression in harvested tumors, total RNA was extracted from xenograft tumor tissue using RNeasy Plus Mini kits (Qiagen) following the manufacturer’s instructions. RNA quantity was determined using a NanoDrop spectrophotometer (Thermo Fisher Scientific). Transcriptional readouts were assessed using a Fluidigm BioMark HD System (Standard BioTools) according to the manufacturer’s recommendations. RNA (100 ng) was subjected to cDNA synthesis and pre-amplification using the High-Capacity cDNA RT Kit and TaqMan PreAmp Master Mix (Thermo Fisher Scientific) per the manufacturer’s protocol. Following amplification, samples were diluted 1:4 with Tris EDTA pH 8.0 and qPCR was conducted using a Fluidigm 96.96 Dynamic Array and the Fluidigm BioMark HD System (Standard BioTools) according to the manufacturer’s recommendations. Cycle threshold (Ct) values were converted to fold changes or percentages in relative expression values (2^-(ddCt)) by subtracting the mean of the housekeeping reference genes from the mean of each target gene followed by subtraction of the mean vehicle dCt from the mean sample dCt.

Blood was harvested from mice treated for 4 days and 3h after the last dosing to quantify the free concentrations of drugs in plasma. Briefly, the concentration of Belvarafenib and Cobimetinib in each sample was determined using a non-validated LC-MS/MS method using labeled internal standards (Cobimetinib: 13C6, Belvarafenib: d5) with qualified curve ranges (Cobimetinib: 1.00 to 100 ng/mL with 2000 ng/mL dilution QC, Belvarafenib: 5.00 to 5000 ng/mL with 75,000 ng/mL dilution QC) using specific columns (Cobimetinib: Waters Xbridge C18, 50 x 2.1 mm, 3.5 um, Belvarafenib: Phenomenex, Onyx Monolithic C18, 50 x 2.0 mm) and MS/MS transition ranges (Cobimetinib: 532.2-249.1, Belvarafenib: 479.1-328.0, 13C6 Cobimetinib: 538.2-255.1, Belvarafenib-d5: 484.1-333.1). The lower limit of quantitation (LLOQ) was 1.00 ng/mL for Cobimetinib and 5.00 ng/mL for Belvarafenib. Free plasma concentrations were calculated by multiplying the plasma concentration in each sample with the fraction unbound in plasma.

### 2.6 Computational dynamic modeling of MAPK signaling

The MARM2 model is written in the PySB framework (https://pysb.org) and describes interactions of the EGFR/MAPK signaling pathway. The model, along with relevant parameters, trained on a range of conditions with MEK and RAF inhibitors, was obtained from Fröhlich, F. and Gerosa, L. et al. [19]. A curation step was performed wherein unnecessary species and their associated model components were removed. The pan-RAF inhibitor Belvarafenib was implemented by setting ep_RAF_RAF_mod_RAFi_double_ddG = 0, removing the reduction in binding affinity of a type 1.5 RAF inhibitor (Vemurafenib) to a partially inhibited RAF dimer [29]. For NRAS Q61 mutants, the hydrolysis rate of NRAS GTP, catalyze_NF1_RAS_gdp_kcatr, was reduced by a factor of 10, and the stability of CRAF dimers, ep_RAF_RAF_mod_RASgtp_double_ddG, was reduced by a factor of 5. Furthermore, since CRAF is the dominant RAF species in NRAS Q61, we removed BRAF in order to greatly reduce model size and computation times. The reduced tendency for phosphorylated CRAF to bind to RAS and form dimers is an important negative feedback mechanism [30,31], which we will refer to as pRAF feedback. To better understand the impacts of this feedback we generate an extra NRAS Q61 model with the feedback removed. Through this process, three models are obtained: BRAF V600E, NRAS Q61 with pRAF feedback, and NRAS Q61 without pRAF feedback.

Each model is converted to a set of ODEs using BNG [32] and then simulated until a steady state is reached. The steady state is achieved when the relative change of all species is less than 0.1% over a period of at least 4 hours. For the steady state dose-responses, 100 inhibitor dose conditions are generated from 10 Cobimetinib doses (0 µM and 9 doses from 10^-2.75^ µM to 10^0^ µM) and 10 Belvarafenib doses (0 µM and 9 doses from 10^-2.25^ µM to 10^0.5^ µM). The initial steady state system is subjected to one of these dose conditions then simulated until the steady state is reached. The full simulation times for all conditions were as follows: BRAF V600E - 475 s, NRAS Q61 without pRAF feedback - 474 s, and NRAS Q61 with pRAF feedback - 330 s (ran on MacBook Pro with M2 Max chip). Bliss values are then generated from the steady state values using the synergy Python library (https://github.com/djwooten/synergy). For the time course responses, the initial steady state system is simulated for 24 hours, then dosed with Cobimetinib (0.5 µM) and either 0 or 133 nM of Belvarafenib. The system is then simulated for 8 additional hours. The full simulation times were as follows: NRAS Q61 without pRAF feedback - 32 s, NRAS Q61 with pRAF feedback - 32 s, and BRAF V600E - 27 s (ran on MacBook Pro with M2 Max chip).

### 2.7 Analysis of drug dose-responses

Cell viability data were processed to relative viability to obtain single-agent fits and metrics (e.g. IC50, Emax and AUC), as well as drug combination fits and metrics such as HSA (Highest Single Agent) and Bliss scores. Briefly, single-agent fits for each drug and cell line were obtained using the *drm* fitting function from the drc R package [33] using a three-parameter (LL.3u) or a four-parameter (LL.4) log-logistic function that relates drug dose to relative viability. For drug combination data, HSA and Bliss scores were calculated as the average of the 10% highest HSA and Bliss excess values observed across the full dose ranges tested, respectively. HSA and Bliss excess values for each dose combination tested were calculated by subtracting the observed response against the expected response under the HSA and Bliss models. As an observed response, we used a smoothened version of the experimental drug combination matrix of relative viability obtained by fitting dose-response curves along every fixed dose of each drug and averaging the fitted values. The HSA expectation matrix was calculated by selecting for each dose combination the maximum response of each individual agent in the observed response. The Bliss expectation was calculated using the Bliss independence formula given as the sum of the responses of the individual drugs minus their product [8]. Data import, processing and calculations were performed using the R package gDR [10].

### 2.8 Projection of in vivo free drug concentrations on in vitro growth responses

Nominal drug concentrations associated with growth viability responses were converted to free drug concentrations in order to project the free drug concentrations measured *in vivo* in mice or patients. Briefly, nominal drug concentrations were multiplied by the fraction unbound (fu) of Belvarafenib and Cobimetinib, which was measured to be 0.034 in 10% FBS media and estimated to be 0.068 in 5% FBS media for Belvarafenib and measured to be 0.196 in 10% FBS media and 0.3 in 5% FBS media for Cobimetinib. To estimate the viability of responses or Bliss excess values at corresponding *in vivo* free drug doses, the matrix with corresponding dose-matrix responses with units converted in free drug concentrations was interpolated using the function *interp2* from the *pracma* R package.

### 2.9 Prediction of tumor growth inhibition in xenografts

GR metric was calculated from the relative viability of IPC-298 cells treated with a combination of Belvarafenib and Cobimetinib by setting an experimentally measured untreated doubling time of 60 hours as described in *Hafner et al.* [16] using the gDR package. The resulting GR metric was converted to control-normalized growth rates, i.e. the growth rate of treated cells divided by the growth rate of control cells. The growth rate of control-treated IPC-298 xenograft tumors was calculated using the doubling time of 18 days estimated from measured tumor volumes to be 0.0385 day^-1^. Using free drug concentrations measured in mice for Belvarafenib and Cobimetinib, corresponding control-normalized growth rates were estimated from the *in vitro* matrix dose-response.

Control-normalized growth rates were multiplied by the baseline tumor growth to predict the growth rate achieved by tumors at any given dosing regime. The obtained growth rates were used in an exponential growth model to simulate tumor volumes in time and compared to experimental data.

### 2.10 Pharmacokinetic (PK) modeling of drug concentrations in patient

Synthetic PK profiles were generated for Belvarafenib and Cobimetinib which recapitulate the population level PK variability expected for each respective compound. For each compound, 500 synthetic PK profiles were generated at each of the following dosing regimens (Belva: 50 mg QD, 100 mg BID, 200 mg BID, and 400 mg BID, Cobi: 20 mg QOD, 20 mg QD, 40 mg QD, and 60 mg QD). These simulations were performed in R 4.1.1 using *mrgsolve* based on the published population PK (popPK) model for Cobimetinib and a popPK model developed on the available individual time-concentration profiles from n=243 patients treated with Belvarafenib in NCT03118817, NCT02405065 and NCT03284502. Both models were developed using the non-linear mixed effects approach as implemented in NONMEM [34]. Simulations were conducted until steady state after which drug levels were recorded for use. In particular, of the 30 days of simulation, days 22–26 were saved for analysis, providing at least two complete cycles of drug concentrations for each condition. Simulated plasma total drug concentrations in ng/mL were divided by the corresponding molecular weight (Belvarafenib = 478.93 g/mol, Cobimetinib= 531.3 g/mol) to obtain total drug concentrations in µM. These were multiplied by the fraction unbound in plasma measured at 0.00258 for Belvarafenib and 0.052 for Cobimetinib.

### 2.11 Clinical tumor growth simulations

A clinical tumor growth inhibition (TGI) model (Claret et al. [35]) was used to describe the tumor dynamics of patients treated in NCT03118817 and NCT03284502. This model was developed using the population approach as implemented in NONMEM version 7.5.0. The model that best described the observed tumor dynamics was a biexponential growth model as described by Stein et al.[36]. In this model, tumor dynamics evolve from an estimated initial tumor size TS_0_, with key treatment-related parameters describing the tumor growth rate constant (KG) (1/week) and tumor shrinkage rate constants (KS) (1/week). Individual empirical Bayesian estimates (EBE) [37] for KG and KS were summarized in melanoma patients and stratified by mutational status. Model-based tumor dynamics were simulated for 1 year for each of these groups based on the mean KG and KS for the group given the same TS_0_ = 50.

## 3. Results

### 3.1. PanRAF and MEK inhibition is additive in BRAF-mutant, but synergistic in NRAS-mutant cell lines

We performed an *in vitro* drug screen to assess the dose-response of 43 melanoma cell lines treated with the type 1.5 “first-generation” RAF inhibitor Vemurafenib [38] and the type 2 “panRAF’’ inhibitor Belvarafenib combined with the allosteric MEK inhibitor Cobimetinib. We measured drug responses using the CellTiter-Glo cell viability assay in a 9-by-9 drug combination matrix design with half-log dilution series starting at the top concentrations of 10 µM for Vemurafenib and Belvarafenib and 5 µM for Cobimetinib. Cell viability readouts were processed using the gDR R package [10] to obtain relative viability and calculate the half-maximal inhibitory concentrations (IC50) (**Figure 1a**) and Bliss scores (**Figure 1b**) as metrics of single-agent potency and combination benefit, respectively. As expected and serving as a control, Vemurafenib as a single agent was found to only inhibit melanoma lines carrying BRAF V600E/K mutations, which signal as BRAF monomers and are thus sensitive to type 1.5 RAF inhibitors that specifically inhibit RAF monomers (**Figure 1a**). In addition to the BRAF V600E/K mutant lines, Belvarafenib also inhibited most melanoma lines with a NRAS hotspot mutation (specifically Q61R, Q61K, Q61V and Q61L) or wild-type for RAS/RAF proteins. This was in line with previous reports [27], as these mutational contexts canonically signal through RAF dimers and are thus sensitive to type 2 RAF inhibitors that block dimeric signaling (**Figure 1a**). The MEK inhibitor Cobimetinib inhibited the growth of most cell lines, validating their broad dependency on MAPK signaling, but interestingly had a much higher potency on cell lines carrying the BRAF V600E/K mutation (log10 mean= -1.66 uM, std= 0.6) than the NRAS mutation (log10 mean= -1.08 uM, std=0.39) or RAS/RAF wild-type (log10 mean= -0.68 uM, std=0.82) (**Figure 1a**).

**Figure 1.**
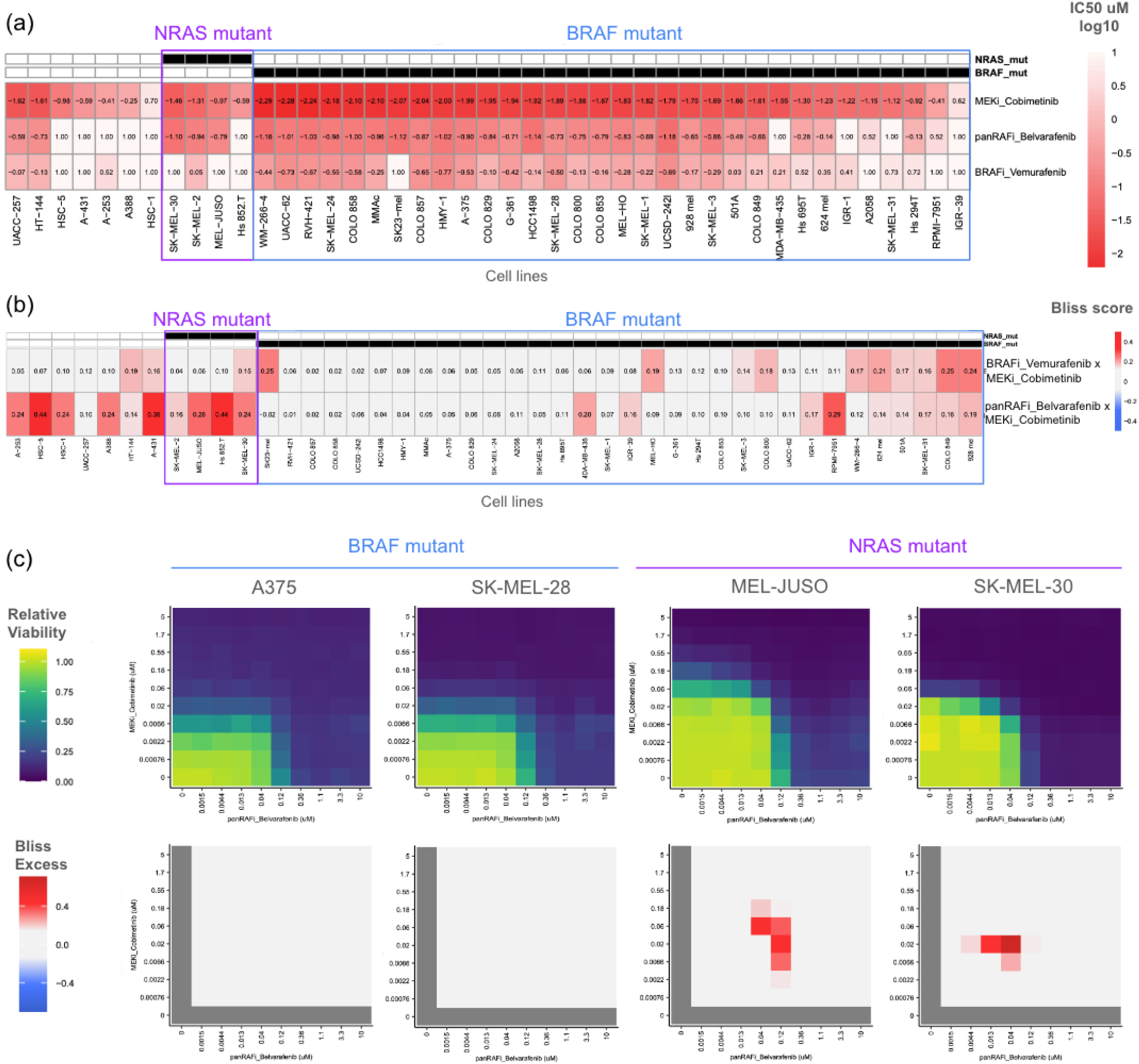
Drug screen reveals additivity of combined pan-RAF and MEK inhibition in BRAF-mutant melanoma, but synergy in NRAS mutant melanoma cell lines. (a) Single agent drug screen on 43 melanoma cell lines including those with BRAF and NRAS mutations. Drug effectiveness quantified via IC50 values. (b) Combination drug screen on the same 43 melanoma cell lines. Drug combination synergies quantified via Bliss scores. (c) Measured in vitro effects of Cobimetinib and Belvarafenib combinations on the relative viability of select cell lines with drug synergies quantified by Bliss excess.

This difference in Cobimetinib’s single-agent potency appeared to extend to the way it combined with Belvarafenib, as quantified by Bliss scores (**Figure 1b**). The combination of Belvarafenib and Cobimetinib presented Bliss scores around zero for most BRAF V600E/K cell lines but positive Bliss scores in most NRAS mutant or RAS/RAF wild-type lines (**Figure 1b**). Bliss scores are calculated as the highest difference between experimentally observed and theoretical expected relative viability based on Bliss independence. With values closer to zero, Bliss scores for BRAF V600E/K melanoma lines (mean=0.10 std=0.06) show that Belvarafenib and Cobimetinib inhibition is mostly additive. High Bliss scores for NRAS mutant (mean=0.27, std=0.12) and RAS/RAF wild-type lines (mean=0.25, std=0.12) highlight a synergistic reduction in relative viability compared to single-agent responses at the same doses. We note that there is a small number of BRAF mutant lines (5/32) that show synergistic pharmacological responses similar to NRAS mutant lines. The dose range at which the maximal benefit is achieved can be visualized by showing relative viability and Bliss excess calculated at each drug dose combination, as shown for representative BRAF and NRAS mutant cell lines (**Figure 1c**). While Bliss excess showed drug additivity across the entire dose-response landscape in BRAF V600E/K lines, NRAS mutant melanoma lines presented a narrow concentration range in which the combination of panRAF and MEK inhibitor highly synergized in inhibiting cancer growth (**Figure 1c**).

### 3.2. Upregulation of MEK phosphorylation in NRAS Q61, but not in BRAF V600 contexts is linked with synergy to panRAF and MEK inhibitors

We reasoned that the different ways in which panRAF and MEK inhibitors combine in NRAS vs BRAF mutant melanomas likely originate from the distinct pathway rewiring caused by these oncogenic mutations. As previously reported, NRAS Q61 signals through RAS-dependent RAF dimers that are sensitive to negative feedback operating on RAFs [31,39] (**Figure 2a**). Instead, BRAF V600E/K signal as RAS-independent RAF monomers that are insensitive to upstream negative feedback (**Figure 2b**).

**Figure 2.**
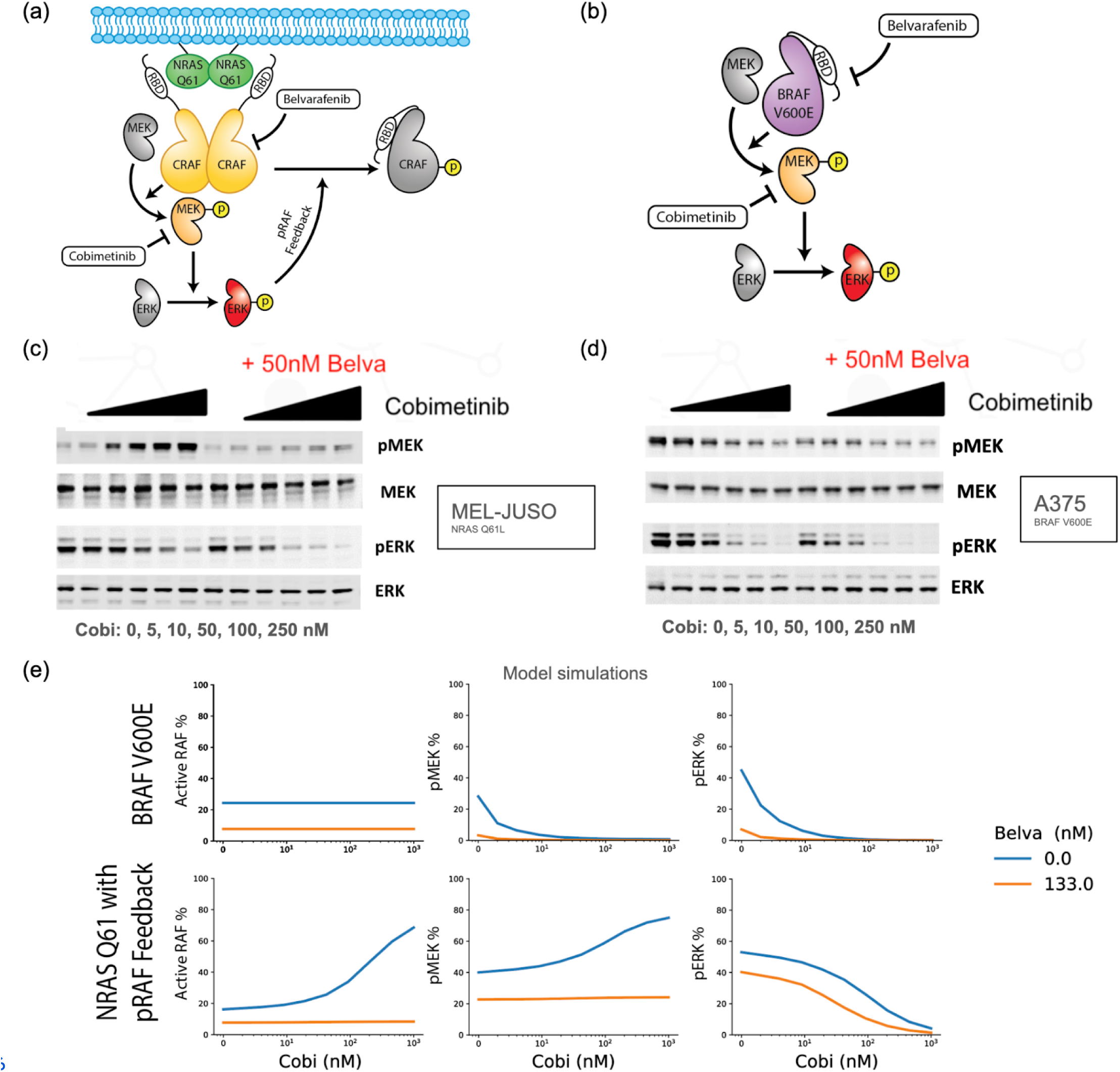
Computational modeling and molecular experiments implicate a negative feedback loop in the response of NRAS vs BRAF mutant melanomas to panRAF and MEK inhibitors. (a-b) Schematic of the MAPK pathway in (a) NRAS Q61 and (b) BRAF V600E melanomas. (c-d) Quantification of pMEK, total MEK, pERK, and total ERK protein levels obtained via western blotting under the indicated combinations of Cobimetinib and Belvarafenib in (c) MEL-JUSO and (d) A-375 cells. (e) Model predictions for steady state percentages of active RAF, pMEK, and pERK under indicated concentrations of Belvarafenib and Cobimetinib. Results are shown for both BRAF V600E and NRAS Q61 models.

To confirm the engagement of negative feedback in NRAS Q61, but not BRAF V600 contexts, we performed western blot experiments with MEL-JUSO (**Figure 2c**) and A-375 cell lines (**Figure 2d**) to measure the phosphorylation status of the MEK and ERK kinases upon inhibition with Cobimetinib, with or without a single dose of Belvarafenib. ERK phosphorylation, the functional output of the MAPK signaling cascade, revealed a trend similar to relative viability readouts: as a single agent, Cobimetinib had lower potency and a shallower dose-response on the NRAS Q61 line MEL-JUSO than the BRAF V600 line A-375 (**Figure 2c-d**). Moreover, combining a fixed dose of Belvarafenib synergized in reducing ERK phosphorylation in the MEL-JUSO lines, but was additive in the A375 line.

MEK phosphorylation measurements were used as a proxy to assess the relief of upstream negative feedback on MAPK signaling. It has previously been shown that upon MEK inhibition, negative feedback release can be observed as a paradoxical increase in pMEK due to higher upstream signaling [40]. Indeed, we found that at doses as low as 10 nM, Cobimetinib induced an increase in MEK phosphorylation in the MEL-JUSO cell line while causing a decrease in the A-375 cell line. Interestingly, the synergy observed between Belvarafenib and Cobimetinib appeared to saturate at the dose of 50 nM Cobimetinib, which corresponds to full engagement of negative feedback as shown by higher MEK phosphorylation in the Cobimetinib single-agent treatment (**Figure 2c-d**). Paradoxical activation of MEK phosphorylation caused by Cobimetinib was abolished by adding Belvarafenib, likely due to a counteracting of the negative feedback relief on RAF dimers (**Figure 2c-d**). These results support the hypothesis that the negative feedback relief observed through pMEK upregulation is linked to the differential response of NRAS Q61 and BRAF V600E lines to Cobimetinib and in combination with Belvarafenib.

### 3.3. Computational model of MAPK signaling implicates negative feedback in the response of NRAS and BRAF mutant melanoma lines to panRAF and MEK inhibitors

To ground this hypothesis on a quantitative framework and disentangle mechanisms of drug synergy, we modified an existing computational model of MAPK signaling that can be instantiated with a BRAF V600 or a NRAS Q61 oncogenic driver [19,20]. Briefly, we implemented and calibrated a previously missing negative feedback that links ERK phosphorylation with an inhibitory phosphorylation of RAF. This phosphorylation reduces the ability for RAF to bind to RAS, dimerize, and facilitate signaling [30,31]. In order to quantitatively assess whether pRAF feedback is capable of explaining the above observations and to better understand the consequences, we made use of the BRAF V600E and NRAS Q61 with pRAF feedback models described in method section 2.6. These models indeed capture the observations made for western blotting data (**Figure 2e**). The NRAS Q61 model exhibits a strong increase in pMEK under single agent Cobimetinib which is significantly diminished with the addition of Belvarafenib while single agent Cobimetinib is effective on the BRAF V600E model. For an NRAS Q61 model with the pRAF feedback removed, there is little to no increase in pMEK in response to Cobimetinib (**Figure S2**) offering support to the hypothesis that negative feedback is key for differential drug responses between NRAS and BRAF mutant tumors.

Next, we used the model to simulate a full drug combination matrix response for Belvarafenib and Cobimetinib. We sampled a dose range focused on the area of synergy and predicted MEK and ERK phosphorylation responses in BRAF V600 and NRAS Q61 contexts (**Figure 3a-b)**. The model predicted that in those dose ranges ERK phosphorylation would be strongly inhibited in the BRAF V600 context by both single agents and in combination. Conversely, it would only strongly inhibit pERK by synergy in the NRAS Q61 context, with a paradoxical activation of pMEK by Cobimetinib. To validate model predictions, we used immunofluorescence-based microscopy to quantify ERK phosphorylation in A-375 and MEL-JUSO cell lines across a 6-by-6 dose dilution matrix of Cobimetinib and Belvarafenib, finding that it accurately and quantitatively matched model predictions (**Figure 3c**). This suggests that the synergistic rather than additive response to panRAF and MEK inhibition observed in NRAS mutant vs BRAF mutant melanoma is driven by the sensitivity to negative feedback of the former compared to the latter. Moreover, drug responses are determined by the degree of inhibition of ERK phosphorylation that is directly translated into cell viability.

**Figure 3.**
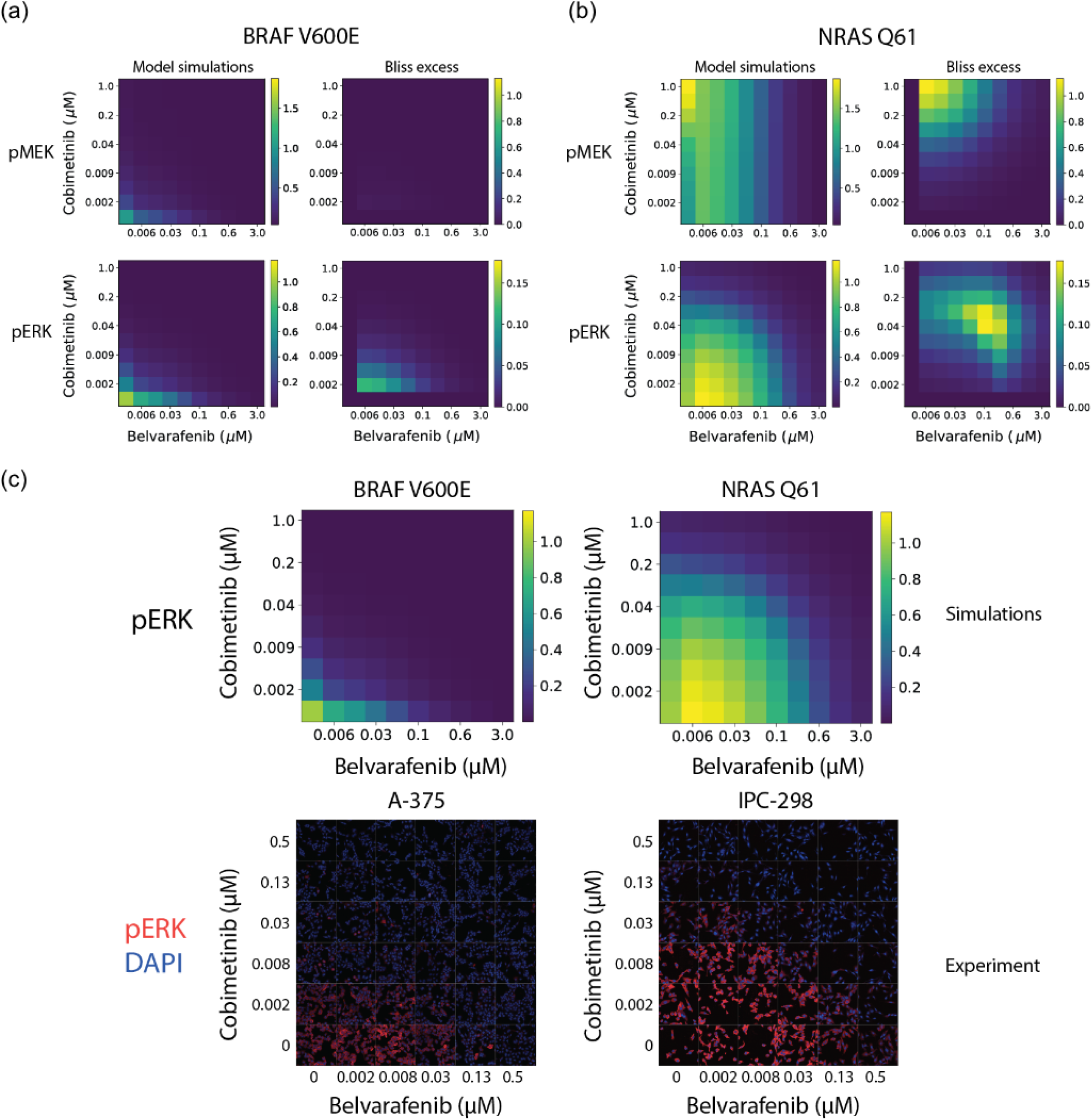
Mechanistic modeling of MAPK signaling quantitatively predicts responses to panRAF and MEK inhibitors in NRAS and BRAF mutant melanoma cell lines. (a-b) Model predictions for pMEK and pERK steady-state levels under indicated concentrations of Belvarafenib and Cobimetinib. Reported values are given relative to drugless conditions. Drug synergy analysis is quantified via excess over Bliss. Values are shown for both (a) BRAF V600E and (b) NRAS Q61 model predictions. (c) Model prediction (top) and immunofluorescence data (bottom) for pERK levels in response to Cobimetinib and Belvarafenib combinations. Values provided for BRAF V600E model and cell line, A-375, (left) and NRAS Q61 model and cell line, IPC-298, (right).

### 3.4. In vitro drug dose-responses assessed at clinically relevant concentrations can accurately predict inhibition of tumor growth in vivo

Next, we wondered if insights obtained from *in vitro* viability responses are relevant to understanding *in vivo* drug dosage and tumor responses. A direct translatability is not obvious as several parameters are different between *in vitro* and *in vivo* settings, such as microenvironment, growth dynamics, cellular states, pharmacokinetic profiles, drug distribution, etc. To directly test translatability, we devised a computational methodology to predict *in vivo* tumor volume responses using as inputs *in vitro* dose-responses and *in vivo* drug concentrations. We applied this methodology to predict tumor responses of IPC-298 melanoma cells grafted in flanks of mice treated for 21 days with clinically relevant doses of Belvarafenib and Cobimetinib (**Figure 4a-b**).

**Figure 4.**
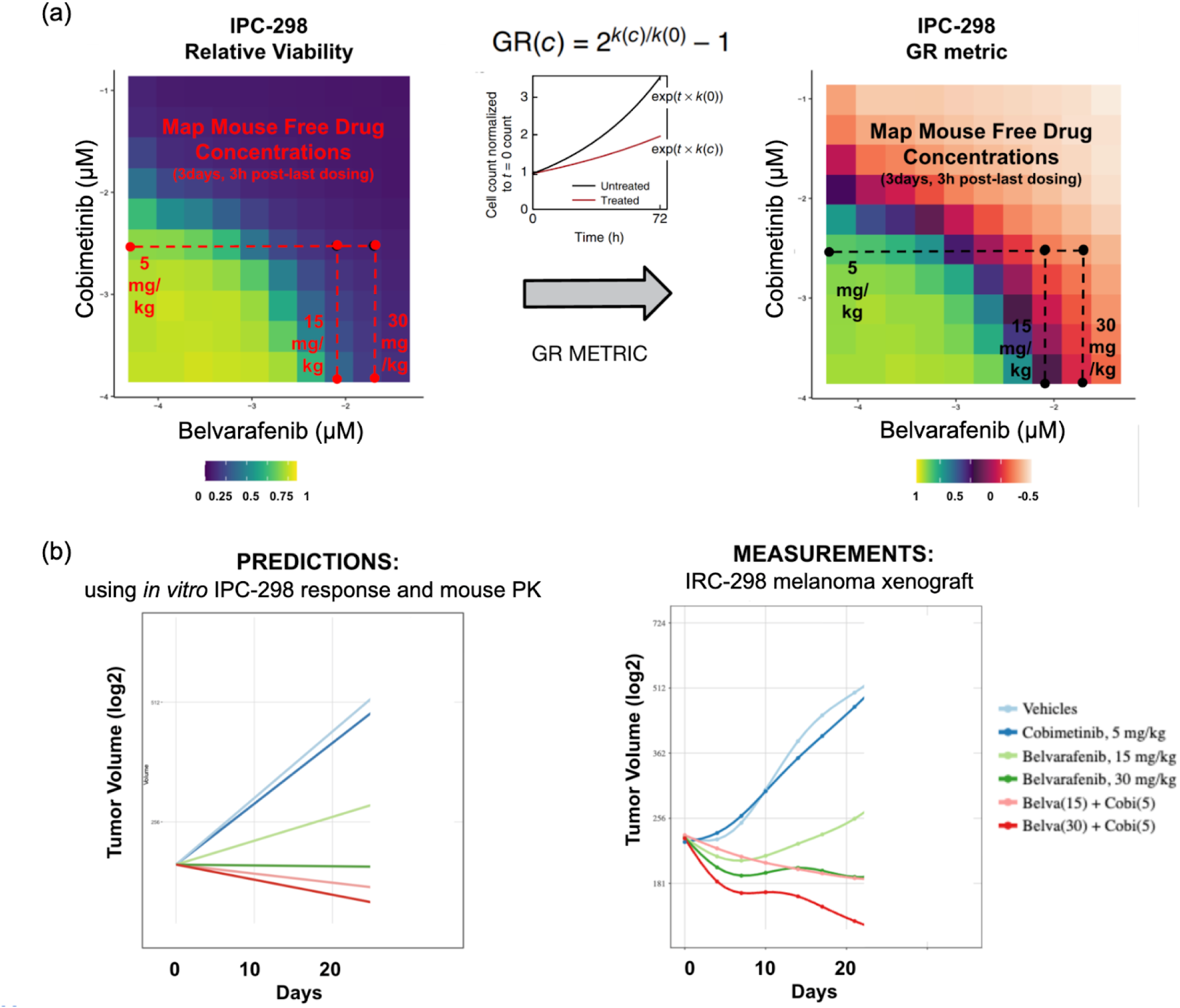
Prediction of *in vivo* xenograft tumor volume control by panRAF and MEK inhibition achieved using *in vitro* cell line response and *in vivo* exposures. (a) Conversion of relative viability to GR metric for IPC-298 in vitro drug responses and projection of mouse PK data onto in vitro responses to obtain predicted tumor growth rates (b) Comparison between predicted tumor growth rates and experimentally measured tumor growth rates. Part of the tumor volume experiments re-analyzed here were previously published in [25].

First, we re-assessed the *in vitro* relative viability of IPC-298 cells using a 10-by-10 dose matrix of Belvarafenib and Cobimetinib with concentration ranges that better match *in vivo* relevant doses (**Figure 4a**). This provides a more refined map on which to score growth inhibition at *in-vivo* drug concentrations compared to the large drug screen. Subsequently, we converted relative viability into growth rate inhibition using the GR metric [15]. Briefly, the baseline doubling rate of IPC-298 cells (60h) was used to back calculate initial seeding cell numbers and calculate the growth rate inhibition at every Belvarafenib and Cobimetinib dose (**Figure 4a**). GR values between one and zero quantify a degree of growth arrest, zero indicates complete stasis and negative values indicate net cell loss (**Figure 4a**). Then, we converted nominal drug concentrations to free drug concentrations by multiplying the fraction unbound (fu) in the serum of each drug (Belvarafenib fu = 0.034, Cobimetinib fu = 0.196).

Second, we projected onto the dose-response matrix the free drug concentrations measured in the plasma of mice treated with 15 mg/kg (free drug = 8 nM) or 30 mg/kg (free drug = 20 nM) of Belvarafenib or 5 mg/kg (free drug = 3 nM) of Cobimetinib QD for 3 days and measured 3 h post last dose. This allowed us to estimate the growth rate inhibition expected from *in vitro* data at the corresponding free drug concentrations for single-agent and combination treatments (**Figure 4a**). Finally, we calculated the baseline growth rate of IPC-298 xenografts in mice treated with vehicle QD for 21 days and scaled the growth rate according to the corresponding *in vitro* growth rate inhibition at each dose regime. This allows us to predict the steady state tumor volume progressions that should be achieved *in vivo* (**Figure 4b**). Comparison with tumor volume growth experimentally measured in mice treated for 21 days showed an accurate prediction of tumor growth dynamics (**Figure 4b**). As single agents, Belvarafenib achieved partial and complete cytostasis at 15 mg/kg and 30 mg/kg, respectively, while Cobimetinib achieved little to no tumor growth inhibition at 5 mg/kg (**Figure 4b**). The addition of 5 mg/kg of Cobimetinib to 15 mg/kg and 30 mg/kg Belvarafenib shifted tumor control from cytostatic to cytotoxic (**Figure 4b**), proving that synergy scored in the *in vitro* setting quantitatively translates into *in vivo* responses. Expression of genes measured at the end of treatment confirmed that improved tumor control is linked to stronger inhibition of genes that report on the activity of MAPK signaling (e.g. FOSL1, DUSP6, SPRY4). Please note that data for three of the five conditions used as comparators for tumor volumes and gene expression analysis here were previously reported in [25]. This confirms the mechanistic basis for synergy previously identified using *in vitro* experiments and computational modeling (**Figure S2b**).

### 3.5. Drug levels required for additive and synergistic responses in NRAS- and BRAF-mutant melanoma can be achieved clinically

We next wondered whether the additive and synergistic behaviors of BRAF and NRAS mutant melanomas observed *in vitro* occur at clinically relevant drug concentrations in patients. In order to evaluate clinically relevant concentrations of Belvarafenib and Cobimetinib, we calculated the average and standard deviation of free drug concentrations from the respective clinical PK models under 16 dose regimens (4 unique dose schemes for each drug) using the simulated responses from day 22 to 26, as described in section 2.10. The average predicted *in vitro* drug combinations were converted to free drug concentrations and projected onto the *in vitro* responses as described in section 2.8 (**Figure 5a**). This approach was used on both the A-375 (BRAF V600E) and IPC-298 (NRAS Q61) cell lines to obtain the GR metric and Bliss excess values for these two mutational contexts at clinically relevant concentrations.

**Figure 5.**
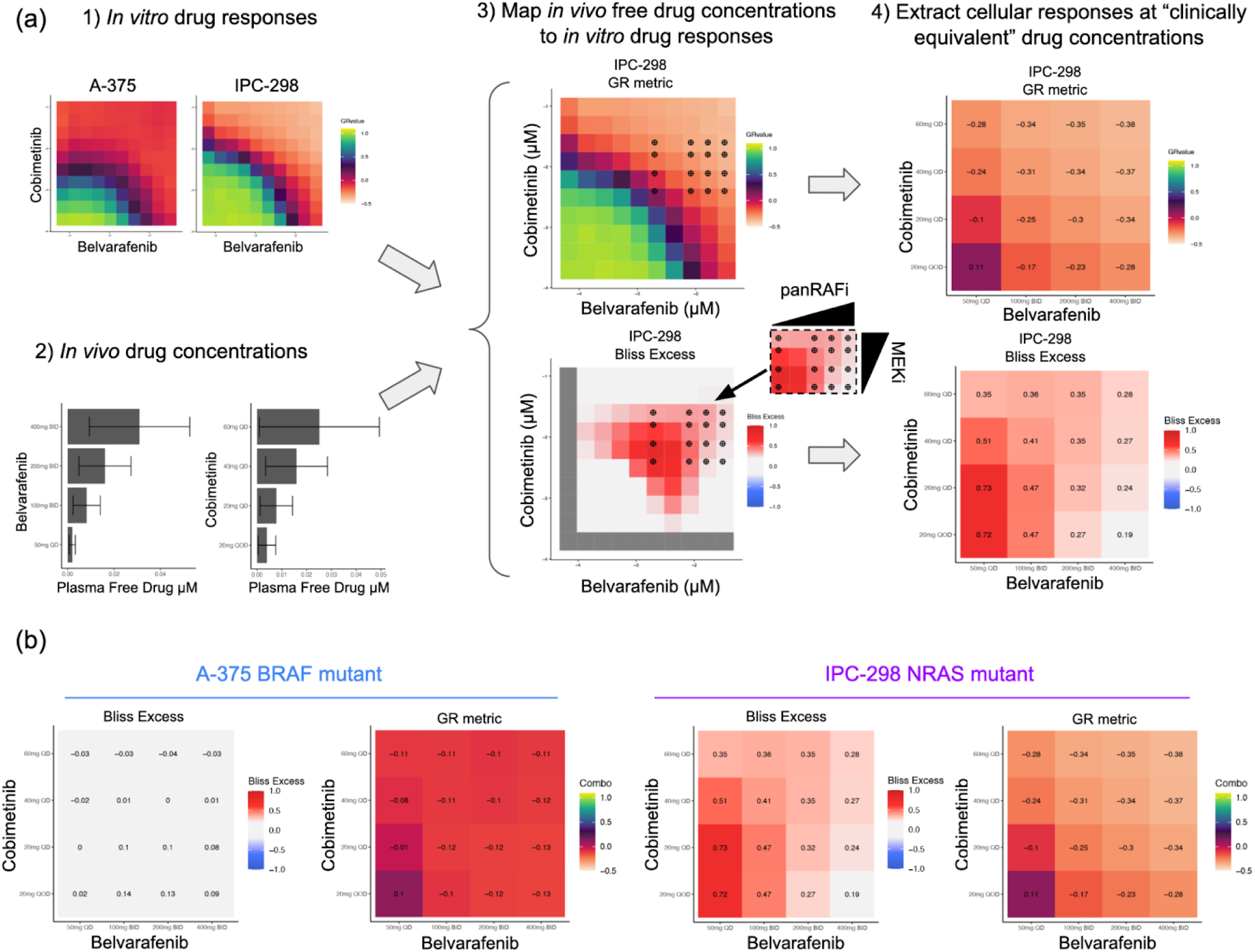
Leveraging synergy in NRAS mutant melanoma at equivalent clinical doses requires at least intermediate MEK inhibition thus allowing lower Belvarafenib doses. (a) Workflow for mapping in vivo free drug concentrations onto in vitro drug responses to predict cell responses and drug synergies at clinically equivalent concentrations. (b) Predicted viability of cell panels and drug synergies at clinically equivalent drug concentrations.

In the BRAF mutant context, all but the weakest clinically realized combinations of Belvarafenib and Cobimetinib perform similarly, inhibiting tumor growth, as shown by the corresponding GR metric values, without significant synergistic effects, as shown by low Bliss excess values (**Figure 5b left**). As a result, we conclude that in BRAF V600E lines there is little motivation to achieve precise drug combination levels in the patient. For these lines, a drug regimen of intermediate intensity should be sufficient to inhibit tumor growth. Conversely, the choice of drug regimen had a greater impact on the extent of growth inhibition in the NRAS mutant context (**Figure 5b right**). Strong tumor inhibition is either achieved with potent Belvarafenib (at 400 mg QD) or Cobimetinib (at 60 mg DQ) single-agent activity or by synergy achieved at intermediate doses, with the highest synergy with good tumor control observed for 100 mg BID Belvarafenib and 20 or 40 mg QD Cobimetinib. This shows that the mutational context creates a different need for dosing of the two combination agents, where leveraging synergy in NRAS mutant melanoma is better achieved at intermediate doses of Cobimetinib that lower the requirement of Belvarafenib to synergize.

As shown by standard deviation errors, we note that the variability in the predicted drug levels is quite large, especially for the higher doses (**Figure 5a bottom left**). This suggests there might be significant issues in achieving a highly synergistic drug combination with precision in individual patients. The NRAS Q61 context thus requires a more thorough analysis of the impacts of this variability to gain insight into which, if any, drug regimens achieve adequate levels of growth inhibition through synergy.

### 3.6. Pharmacokinetic variability in patients highlights precision requirement for synergistic responses in NRAS mutant melanoma tumors

We decided to assess the role that the patient-to-patient variability in pharmacokinetic profiles has in leveraging synergistic vs additive responses. The PK models in this study provide drug levels for individual, virtual patients, which enables us to develop a mapping from each single patient’s PK profile to a distribution of drug effects the patient experiences, i.e. GR metric and Bliss Excess values. To accomplish this, we obtain the patient’s free drug concentrations once per hour over the course of 48 hours (**Figure S3**), then project these concentrations onto the GR metric and Bliss scores of each mutational context in the same way we projected the average free drug levels in section 3.5. Doing this for multiple patients reveals the impacts of patient-to-patient variation as well as the effects resulting from the temporal variation of drug levels (**Figure 6a**). From this we see a single drug regimen can generate different responses within a population.

**Figure 6.**
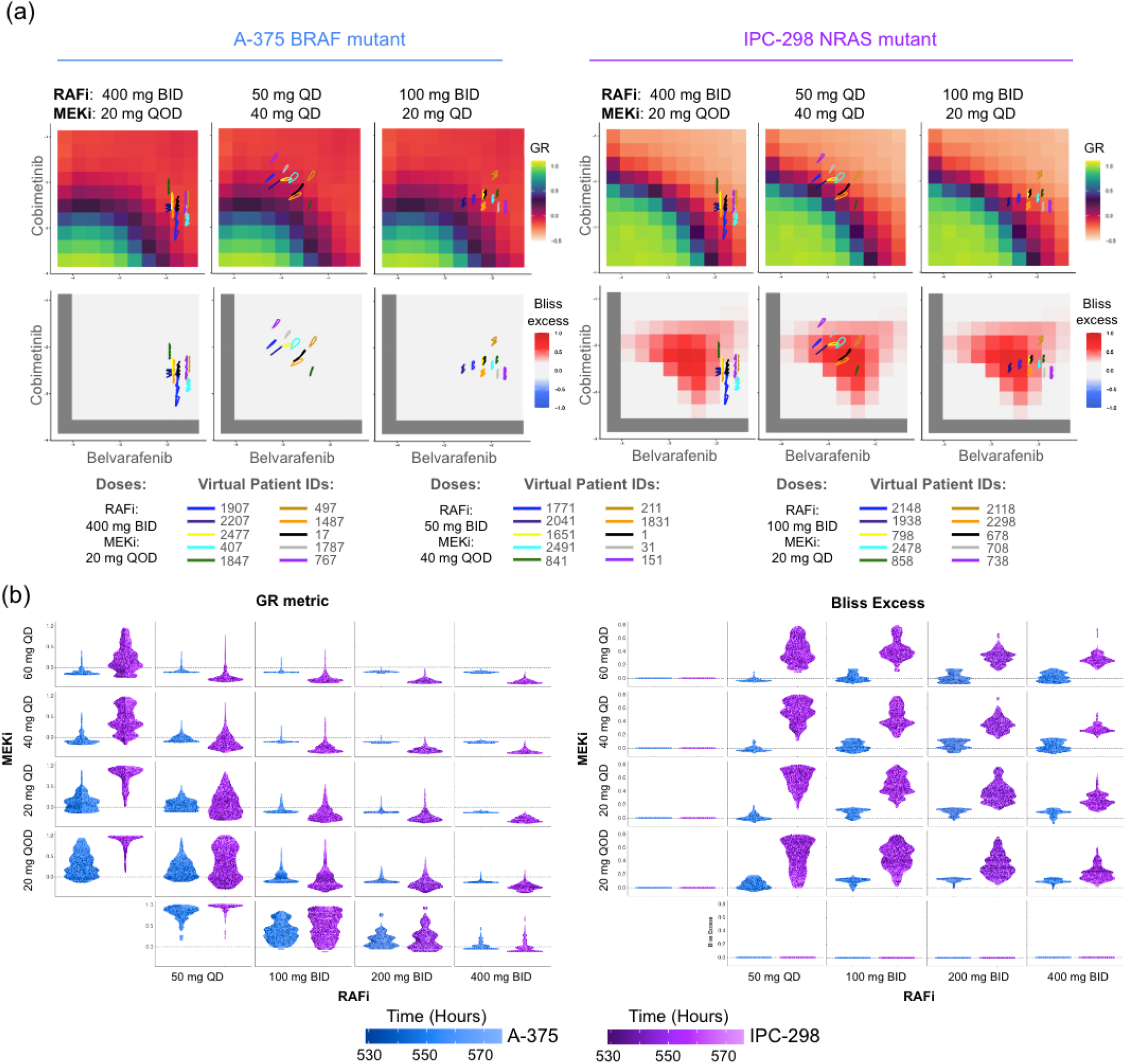
Pharmacokinetic variability in patients limits the precision to obtain synergistic responses in NRAS mutant melanoma tumors (a) Individual virtual patient PK trajectories resulting from the indicated drug regimen projected onto in vitro responses. (b) The distribution of GR metric values (left) and Bliss excess values (right) measured from 75 single patient trajectories. Multiple drug regimens are compared, rows and columns labels indicate the Cobimetinib and Belvarafenib doses used in the specific drug regimen.

This indicates a significant challenge for treatments; a given regimen might, for example, work well for one patient, but have less effect for another. In order to gain a better understanding of which regimens consistently result in high benefit/low tumor growth across all patients and times, we examine the full distribution of predicted effects that result from a given drug regimen (**Figure 6b**). We find that because BRAF V600E tumors lack significant synergy (low Bliss excess) and still achieve consistently strong tumor suppression (high GR values) from drug regimens with as low dosing as Belvarafenib 100 mg QD and Cobimetinib 20 mg QD. Therefore, we conclude that drug additivity imposes no strict requirements on the precision of dosing in this mutational context.

Conversely, NRAS Q61 tumors are seen to achieve tumor control by significant synergy (high Bliss excess) for drug combinations that leverage partial single-agent MEK inhibition at 20 and, even better, 40 mg QD regimen, which combine well with doses as low as 50 mg QD and 100 mg BID of Belvarafenib. Distribution of growth inhibition measured by GR and synergy by Bliss excess visualized via violin plots show, however, that combinations with 50 mg QD Belvarafenib suffer from incomplete responses due to the large variability in free drug concentrations in individual patients. This happens because combinations with 50 mg QD Belvarafenib lie very close to the synergy boundary in the dose landscape and fluctuations bring the response outside of the synergistic regimes (**Figure 6a-b**). Of all the synergistic combinations, Cobimetinib at 40 mg QD with Belvarafenib at 100 mg BID seem to achieve consistent tumor control with lower patient-to-patient variability and moderate single agent-activities, thus representing an ideal drug-sparing synergistic point in the dose landscape. This underscores the importance of using dose regimes with high synergy when treating NRAS Q61 tumors to achieve strong effects while minimizing the effect of pharmacokinetic fluctuations.

### 3.7. Clinical trials support distinct combinability of panRAF and MEK inhibitors in BRAF and NRAS mutant patients

To ascertain the validity of insights from modeling and experiments, we analyzed limited data available from Phase 1 clinical trials combining Belvarafenib and Cobimetinib in the treatment of melanoma patients. We fit a clinical tumor growth inhibition (TGI) model [35] to describe the tumor dynamics of patients treated in clinical trials NCT03118817 and NCT03284502, as described in section 2.11. The model describes the observed tumor dynamics with a biexponential growth model with tumor dynamics evolving for one year from the estimated initial tumor size, with tumor growth rate and tumor shrinkage rate constants summarized in melanoma patients and stratified by mutational status [36]. The simulations provide support for the differential contribution of increasing the Cobimetinib dose in the BRAF-mutant vs NRAS-mutant setting. As we predicted, the supralinear impact on growth from increasing Cobimetinib doses on the NRAS mutant tumors subjected to a constant Belvarafenib dose (**Figure 7b bottom**) indicates the presence of synergistic effects. While the more linear impact on growth from increasing Cobimetinib doses on the BRAF mutant tumor subjected to a constant Belvarafenib dose (**Figure 7b top**) indicates the drugs are acting in a more additive fashion. This synergy does appear important for reaching desired effects in NRAS mutant tumors, with a combination of Cobimetinib and Belvarafenib outperforming single agent Belvarafenib at suppressing tumor growth.

**Figure 7.**
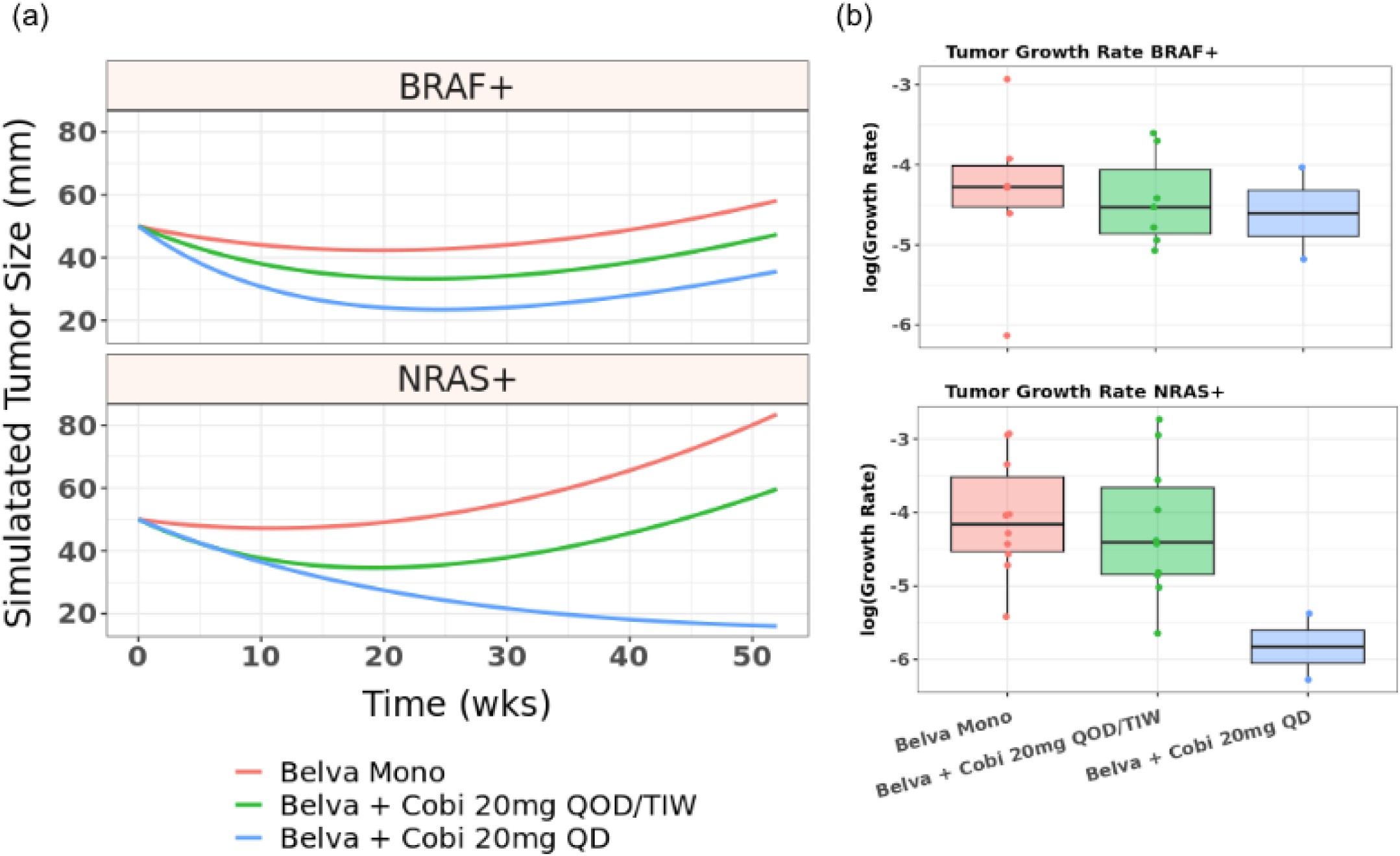
Tumor growth inhibition in patients simulated using a model trained on Phase 1 clinical trials support the additive vs synergistic dose landscape of BRAF vs NRAS mutant melanoma patients for panRAF and MEK co-inhibition (a) Simulated tumor growth under indicated Belvarafenib and Cobimetinib regimens for BRAF and NRAS mutant melanoma patients. Belva mono is 400/450 mg BID (b) Distribution of tumor growth rates for indicated drug regimen within simulated populations of patients with BRAF V600E (top) or NRAS Q61 (bottom) tumors.

Clinical data allow us to assess another key information for the design of drug combinations not included in our analysis, namely if tolerability is a relevant issue that constrains drug regimens. In the clinical trial NCT03284502, the regimen of Belvarafenib 200 mg BID continuously and Cobimetinib 40 mg QD 21/7 led to 3 dose-limiting toxicities (DLTs) (G3 colitis, G3 diarrhea, G3 nausea) in 2 patients [41]. These and other reported treatment-emergent toxicities (“dermatitis acneiform, diarrhea, constipation, and increase in blood creatine phosphokinase”) suggest on-target toxicity on wild-type MAPK signaling. Consequently, Cobimetinib was reduced to 20 mg QD while Belvarafenib was dose escalated to 300 mg BID, which did not result in DLTs [41]. Our analysis described in **Figure 6b** shows that at 200 mg BID Belvarafenib and 40 mg QD Cobimetinib, Belvarafenib and Cobimetinib are already both substantially active as single agents in NRAS mutant cells, suggesting that the combination is not leveraging synergy as effectively and is likely impinging on wild-type MAPK signaling. Increasing Belvarafenib to 300 mg BID while reducing Cobimetinib to 20 mg QD shifts the contribution to mostly Belvarafenib as single-agent, likely reducing toxicity but also losing synergistic effects on NRAS-mutant tumors. Our analysis suggests maintaining Cobimetinib at 40 mg QD or QOD while reducing Belvarafenib to as low as 50-100 mg QD/BID is an alternative approach to de-escalate dose intensity which might better leverage synergy of tumor inhibition without invoking strong single-agent effects, the possible culprits of toxicity. To the best of our knowledge, this regime of intermediate Cobimetinib dose and low Belvarafenib dose remains untested in the clinic.

## 4. Discussion

This study integrates drug response*s,* signaling modeling and pharmacokinetic simulations to identify mutational scenarios sensitive to specific co-dosing regimens in precision therapy for melanoma. Our main finding is that panRAF and MEK inhibition exhibit additive effects in BRAF-mutant tumors and synergistic effects in NRAS-mutant tumors and that this difference translates into distinct requirements in terms of dosing regimens and dosing precision in the clinic. Our approach addresses a number of shortcomings typically encountered in translating *in vitro* to *in vivo* drug responses. In the following, we will elaborate on these findings as well as discuss the constraints and limitations of our own methodology.

We identified differences in the benefit of panRAF and MEK co-inhibition through a drug screen of 43 melanoma cell lines. While the screen was strongly biased for BRAF V600 mutations, high synergy was evident in four NRAS mutant lines as quantified by Bliss excess analysis. Our analysis extended beyond these traditional combination metrics by projecting *in vivo* drug doses onto drug combination responses. Key to this projection was gathering information on free drug concentrations coming from *in vivo* xenograft experiments and pharmacokinetic models trained using clinical data. Our approach confirmed that the additivity and synergy detected in vitro apply at clinically achievable doses of Belvarafenib and Cobimetinib. The computational tool we developed for this analysis aids in the definition of dose-response matrices reflective of clinical conditions and is publicly available to encourage use in the scientific community.

An issue with projecting clinical concentrations on drug dose-response data is the translatability from *in vitro* to *in vivo*. We found that converting relative viability to growth rate inhibition via GR metric allowed for precise prediction of tumor inhibition in a xenograft model. This methodology was previously shown to be effective for single agent drugs, but with the necessity of an inferred conversion factor to relate *in vitro* and *in vivo* drug concentrations [16]. We found that in our system this factor is unnecessary, i.e. it is unity. It is possible that other drug combinations or cell lines will not enjoy this direct correspondence. Using the approach we develop here to systematically assess conversion factors across drug combinations and cancer models should help extract the principles by which *in vitro* responses translate to *in vivo* settings, guiding translatability of pre-clinical studies. While our findings suggest this is possible, a notable limitation is the reliance on cell lines and xenografts, which might not accurately represent clinical response as they may not fully encapsulate the intricate biology of patient tumors and lack critical elements such as the immune system.

Mechanistically, we identified a negative feedback on RAF dimers in NRAS mutant melanoma as the likely culprit behind their lower sensitivity to single-agent MEK inhibition and synergistic response to panRAF co-inhibition. These findings largely confirm prior research [26–28], but were extended using computational modeling of signal transduction to provide a quantitative framework for understanding and predicting mechanisms of drug adaptation. We have shown that a previously developed model of MAPK signaling [19,20] could be extended to explore synergy mechanisms specific to these mutational contexts. Moreover, we used the model to design experiments that validated the key link between the degree of ERK inhibition achieved in BRAF and NRAS mutant cell lines and drug responses. As noted in the results section, there was a small fraction of BRAF-mutant lines that exhibited synergistic responses similar to NRAS-mutant lines. Mechanistic insights from modeling indicate that these BRAF-mutant lines might activate dimeric RAF signaling either at baseline or in response to treatment, therefore suggesting that drug synergy might also be required to curb resistance mechanisms in BRAF-mutant tumors.

With a mechanistic understanding in hand, next we assessed drug responses at clinical-relevant concentrations to retrospectively evaluate dosing regimes tested in the clinic and foresight alternative strategies. As scored through the lenses of pre-clinical data, we realized that the initial combination tested in the clinic of 200 mg BID Belvarafenib and 40 mg QD Cobimetinib does not fully leverage synergy since both drugs, but especially Belvarafenib, are quite effective as single-agents. Interestingly, this dose regime was also not well tolerated in the clinic, most likely due to on-target toxicity. Our analysis suggests that to fully leverage synergy and reduce single-agent activity, Cobimetinib could be kept at 40 mg QD or QOD dosing while Belvarafenib could be reduced substantially to 50 or 100 mg QD/BID. This strategy agrees with the preclinical evidence that synergy is best leveraged when the negative feedback elicited by MEK inhibition is partially active to dramatically potentiate panRAF inhibition. We hypothesize that the alternative strategy of lowering Cobimetinib to 20 mg QD and escalating Belvarafenib to 300 mg BID might come at the cost of losing single-agent potency of MEK inhibition and, therefore, drug synergy.

Although supported by preclinical data for efficacy, utilizing drug synergy in a regimen of intermediate MEK inhibition and low panRAF inhibition to minimize on-target toxicities in the clinic remains to be validated. In principle, on-target toxicities could be reduced if the mechanisms behind drug synergy are not strongly operating in healthy tissues. While evidence seems to suggest the tendency of synergy in therapeutic effects to be significantly stronger than the synergy in toxic effects [43], the picture is far from clear. On the downside, the negative feedback mechanisms behind drug synergy operating on Ras signaling and RAF dimerization are presumably active in healthy cells. In contrast, NRAS Q61 hotspot mutations disrupt the Ras loading cycle in a manner that likely amplifies dependency on negative feedback—and thus drug synergy—relative to healthy cells. A lack of cell line models that accurately represent signaling in normal tissue complicates experimental verification of these hypotheses. Estimating the therapeutic window of wild-type versus mutant signaling presents a promising direction for dynamic signaling modeling, especially when parameters for wild-type signaling are quantifiable. Ultimately, the clinical experiment of a regimen involving intermediate MEK inhibition and low panRAF inhibition needs to be implemented to assess the effect on toxicity. A main result of this work is to show that this regime remains so far likely untested for Belvarafenib and Cobimetinib.

An insight revealed by analyzing patient-to-patient pharmacokinetic variability is the degree of precision in dosing needed to leverage synergy in the clinics. We have observed that the synergistic space of the dose landscape is pretty narrow compared to the fluctuations in free drug concentrations across patients. For each regime in which average drug concentrations were solidly in the synergistic space, we found some patients whose fluctuations in drug levels positioned them outside of synergy. This has implications for how preclinical evidence of synergy should be applied to implement drug combinations in the clinic. Conceptually, our observations might propose a more general principle often overlooked in clinical development. Using clinical data on patient responses, it has been shown that most drug combinations in the clinic in practice act independently or additively, even when pre-clinical work suggested strong synergy [42]. We find it unlikely that mechanisms of synergy identified pre-clinically do not operate in human tumors. Here, we argue that synergy might not often be observed clinically because of the practical issue of maintaining drug concentrations within the synergistic regimes. We suggest that the methodology developed here can be applied early on in clinical decision making to inform on the likelihood of achieving and maintaining synergy in a patient population.

## 5. Conclusions

In this study, we explore the use of preclinical cell line drug response data alongside computational modeling to determine the optimal dosages of pan-RAF (Belvarafenib) and MEK (Cobimetinib) inhibitors for melanoma treatment. The main findings is that the two main oncogenic drivers in melanoma, BRAF V600 and NRAS Q61 hotspot mutations, result in different underlying signaling biology requiring different treatment regimes using the same drugs. We show that most combinatorial dose regimens achievable in the clinic are effective for treating BRAF-mutant melanoma thanks to single-agent higher potency and drug additivity, whereas NRAS-mutant melanoma requires more precise dosing to harness drug synergy, posing practical implementation challenges due to interpatient pharmacokinetic variability.

Our research underscores that precision medicine should not only aim to identify the most effective drug combination for a given indication, but also tailor dosing regimens to match the pathway biology driven by mutational mechanisms, among other biologic factors. In these contexts, the need for precision dosing becomes imperative, demanding thorough examination within both pre-clinical and translational research frameworks. By introducing a novel methodological approach, our study seeks to tackle the challenges associated with implementing precision dosing strategies, propelling the efforts to enhance the personalization of cancer treatment.

## Author Contributions

Conceptualization, L.G.; methodology, A.G., F.S, B.C., E.L., R.M, D.D.L, T.H. and L.G.; software, A.G. and L.G; validation, A.G., F.S, E.L., R.M, D.D.L and L.G.; formal analysis, A.G. and L.G.; investigation, A.G, F.S., S.F. and L.G..; resources, U.S., S.F, S.M., P.D. and L.G.; data curation, A.G. and L.G..; writing—original draft preparation, A.G and L.G.; writing—review and editing, A.G., S.A.F and L.G.; visualization, A.G. and L.G..; supervision, L.G.; project administration, L.G..; funding acquisition, U.S., S.F, S.M., P.D. and L.G. All authors have read and agreed to the published version of the manuscript.

## Institutional Review Board Statement

Genentech is an AAALAC-accredited facility and all animal activities in the research studies were conducted under protocols approved by the Genentech Institutional Animal Care and Use Committee (IACUC).

## Informed Consent Statement

All studies were performed in accordance with the Declaration of Helsinki and participants provided written informed consent. Additional details of the studies and a complete list of inclusion and exclusion criteria can be found on clinicaltrials.gov (NCT03118817, NCT02405065 and NCT03284502).

## Data Availability Statement

Experimental data in machine readable formats and R/Python code to reproduce the computational analysis in this work are available at GitHub: https://github.com/lgerosa/panRAFi_MEKi_combo.

## Acknowledgments

The authors thank Marc Hafner, Cassie Chou, Yibing Yan, Jennifer Eng-Wong and Michael Dolton at Genentech/Roche for helpful discussions.

## References

1. Behan, F.M.; Iorio, F.; Picco, G.; Gonçalves, E.; Beaver, C.M.; Migliardi, G.; Santos, R.; Rao, Y.; Sassi, F.; Pinnelli, M.;, et al. Prioritization of Cancer Therapeutic Targets Using CRISPR–Cas9 Screens. Nature 2019, 568, 511–516, doi:10.1038/s41586-019-1103-9.

2. Tsherniak, A.; Vazquez, F.; Montgomery, P.G.; Weir, B.A.; Kryukov, G.; Cowley, G.S.; Gill, S.; Harrington, W.F.; Pantel, S.; Krill-Burger, J.M.;, et al. Defining a Cancer Dependency Map. Cell 2017, 170, 564–576.e16, doi:10.1016/j.cell.2017.06.010.

3. Pagliarini, R.; Shao, W.; Sellers, W.R. Oncogene Addiction: Pathways of Therapeutic Response, Resistance, and Road Maps toward a Cure. EMBO Rep. 2015, 16, 280–296, doi:10.15252/embr.201439949.

4. Kolch, W.; Halasz, M.; Granovskaya, M.; Kholodenko, B.N. The Dynamic Control of Signal Transduction Networks in Cancer Cells. Nat. Rev. Cancer 2015, 15, 515–527, doi:10.1038/nrc3983.

5. Labrie, M.; Brugge, J.S.; Mills, G.B.; Zervantonakis, I.K. Therapy Resistance: Opportunities Created by Adaptive Responses to Targeted Therapies in Cancer. Nat. Rev. Cancer 2022, 22, 323–339, doi:10.1038/s41568-022-00454-5.

6. Bashi, A.C.; Coker, E.A.; Bulusu, K.C.; Jaaks, P.; Crafter, C.; Lightfoot, H.; Milo, M.; McCarten, K.; Jenkins, D.F.; Van Der Meer, D.;, et al. Large-Scale Pan-Cancer Cell Line Screening Identifies Actionable and Effective Drug Combinations. Cancer Discov. 2024, 14, 846–865, doi:10.1158/2159-8290.CD-23-0388.

7. Jaaks, P.; Coker, E.A.; Vis, D.J.; Edwards, O.; Carpenter, E.F.; Leto, S.M.; Dwane, L.; Sassi, F.; Lightfoot, H.; Barthorpe, S.;, et al. Effective Drug Combinations in Breast, Colon and Pancreatic Cancer Cells. Nature 2022, 603, 166–173, doi:10.1038/s41586-022-04437-2.

8. Vlot, A.H.C.; Aniceto, N.; Menden, M.P.; Ulrich-Merzenich, G.; Bender, A. Applying Synergy Metrics to Combination Screening Data: Agreements, Disagreements and Pitfalls. Drug Discov. Today 2019, 24, 2286–2298, doi:10.1016/j.drudis.2019.09.002.

9. Wooten, D.J.; Meyer, C.T.; Lubbock, A.L.R.; Quaranta, V.; Lopez, C.F. MuSyC Is a Consensus Framework That Unifies Multi-Drug Synergy Metrics for Combinatorial Drug Discovery. Nat. Commun. 2021, 12, 4607, doi:10.1038/s41467-021-24789-z.

10. Vuong, Allison; Czech, Bartosz; Gladki, Arkadiusz; Hafner, Marc; Scigocki, Dariusz; Smola, Janina; Mocanu, Sergiu gDR: Umbrella Package for R Packages in the gDR Suite 2023.

11. Mammoliti, A.; Smirnov, P.; Safikhani, Z.; Ba-Alawi, W.; Haibe-Kains, B. Creating Reproducible Pharmacogenomic Analysis Pipelines. Sci. Data 2019, 6, 166, doi:10.1038/s41597-019-0174-7.

12. Summerfield, S.G.; Yates, J.W.T.; Fairman, D.A. Free Drug Theory – No Longer Just a Hypothesis? Pharm. Res. 2022, 39, 213–222, doi:10.1007/s11095-022-03172-7.

13. Clarke, M.A.; Fisher, J. Executable Cancer Models: Successes and Challenges. Nat. Rev. Cancer 2020, 20, 343–354, doi:10.1038/s41568-020-0258-x.

14. Adam, G.; Rampášek, L.; Safikhani, Z.; Smirnov, P.; Haibe-Kains, B.; Goldenberg, A. Machine Learning Approaches to Drug Response Prediction: Challenges and Recent Progress. *Npj Precis*. Oncol. 2020, 4, 19, doi:10.1038/s41698-020-0122-1.

15. Diegmiller, R.; Salphati, L.; Alicke, B.; Wilson, T.R.; Stout, T.J.; Hafner, M. Growth-rate Model Predicts in Vivo Tumor Response from in Vitro Data. CPT Pharmacomet. Syst. Pharmacol. 2022, 11, 1183–1193, doi:10.1002/psp4.12836.

16. Hafner, M.; Niepel, M.; Chung, M.; Sorger, P.K. Growth Rate Inhibition Metrics Correct for Confounders in Measuring Sensitivity to Cancer Drugs. Nat. Methods 2016, 13, 521–527, doi:10.1038/nmeth.3853.

17. Pillai, M.; Hojel, E.; Jolly, M.K.; Goyal, Y. Unraveling Non-Genetic Heterogeneity in Cancer with Dynamical Models and Computational Tools. Nat. Comput. Sci. 2023, 3, 301–313, doi:10.1038/s43588-023-00427-0.

18. McFall, T.; Diedrich, J.K.; Mengistu, M.; Littlechild, S.L.; Paskvan, K.V.; Sisk-Hackworth, L.; Moresco, J.J.; Shaw, A.S.; Stites, E.C. A Systems Mechanism for KRAS Mutant Allele–Specific Responses to Targeted Therapy. Sci. Signal. 2019, 12, eaaw8288, doi:10.1126/scisignal.aaw8288.

19. Fröhlich, F.; Gerosa, L.; Muhlich, J.; Sorger, P.K. Mechanistic Model of MAPK Signaling Reveals How Allostery and Rewiring Contribute to Drug Resistance. Mol. Syst. Biol. 2023, 19, e10988, doi:10.15252/msb.202210988.

20. Gerosa, L.; Chidley, C.; Fröhlich, F.; Sanchez, G.; Lim, S.K.; Muhlich, J.; Chen, J.-Y.; Vallabhaneni, S.; Baker, G.J.; Schapiro, D.;, et al. Receptor-Driven ERK Pulses Reconfigure MAPK Signaling and Enable Persistence of Drug-Adapted BRAF-Mutant Melanoma Cells. Cell Syst. 2020, 11, 478–494.e9, doi:10.1016/j.cels.2020.10.002.

21. Rukhlenko, O.S.; Khorsand, F.; Krstic, A.; Rozanc, J.; Alexopoulos, L.G.; Rauch, N.; Erickson, K.E.; Hlavacek, W.S.; Posner, R.G.; Gómez-Coca, S.;, et al. Dissecting RAF Inhibitor Resistance by Structure-Based Modeling Reveals Ways to Overcome Oncogenic RAS Signaling. Cell Syst. 2018, 7, 161–179.e14, doi:10.1016/j.cels.2018.06.002.

22. Stites, E.C.; Shaw, A.S. Quantitative Systems Pharmacology Analysis of KRAS G12C Covalent Inhibitors. CPT Pharmacomet. Syst. Pharmacol. 2018, 7, 342–351, doi:10.1002/psp4.12291.

23. Chapman, P.B.; Hauschild, A.; Robert, C.; Haanen, J.B.; Ascierto, P.; Larkin, J.; Dummer, R.; Garbe, C.; Testori, A.; Maio, M.;, et al. Improved Survival with Vemurafenib in Melanoma with BRAF V600E Mutation. N. Engl. J. Med. 2011, 364, 2507–2516, doi:10.1056/NEJMoa1103782.

24. Chapman, P.B.; Solit, D.B.; Rosen, N. Combination of RAF and MEK Inhibition for the Treatment of BRAF-Mutated Melanoma: Feedback Is Not Encouraged. Cancer Cell 2014, 26, 603–604, doi:10.1016/j.ccell.2014.10.017.

25. Yen, I.; Shanahan, F.; Lee, J.; Hong, Y.S.; Shin, S.J.; Moore, A.R.; Sudhamsu, J.; Chang, M.T.; Bae, I.; Dela Cruz, D.;, et al. ARAF Mutations Confer Resistance to the RAF Inhibitor Belvarafenib in Melanoma. Nature 2021, 594, 418–423, doi:10.1038/s41586-021-03515-1.

26. Yuan, X.; Tang, Z.; Du, R.; Yao, Z.; Cheung, S.; Zhang, X.; Wei, J.; Zhao, Y.; Du, Y.; Liu, Y.;, et al. RAF Dimer Inhibition Enhances the Antitumor Activity of MEK Inhibitors in *K-RAS* Mutant Tumors. Mol. Oncol. 2020, 14, 1833–1849, doi:10.1002/1878-0261.12698.

27. Yen, I.; Shanahan, F.; Merchant, M.; Orr, C.; Hunsaker, T.; Durk, M.; La, H.; Zhang, X.; Martin, S.E.; Lin, E.;, et al. Pharmacological Induction of RAS-GTP Confers RAF Inhibitor Sensitivity in KRAS Mutant Tumors. Cancer Cell 2018, 34, 611–625.e7, doi:10.1016/j.ccell.2018.09.002.

28. Whittaker, S.R.; Cowley, G.S.; Wagner, S.; Luo, F.; Root, D.E.; Garraway, L.A. Combined Pan-RAF and MEK Inhibition Overcomes Multiple Resistance Mechanisms to Selective RAF Inhibitors. Mol. Cancer Ther. 2015, 14, 2700–2711, doi:10.1158/1535-7163.MCT-15-0136-T.

29. Cook, F.A.; Cook, S.J. Inhibition of RAF Dimers: It Takes Two to Tango. Biochem. Soc. Trans. 2021, 49, 237–251, doi:10.1042/BST20200485.

30. Dougherty, M.K.; Müller, J.; Ritt, D.A.; Zhou, M.; Zhou, X.Z.; Copeland, T.D.; Conrads, T.P.; Veenstra, T.D.; Lu, K.P.; Morrison, D.K. Regulation of Raf-1 by Direct Feedback Phosphorylation. Mol. Cell 2005, 17, 215–224, doi:10.1016/j.molcel.2004.11.055.

31. Yu, A.; Nguyen, D.H.; Nguyen, T.J.; Wang, Z. A Novel Phosphorylation Site Involved in Dissociating RAF Kinase from the Scaffolding Protein 14-3-3 and Disrupting RAF Dimerization. J. Biol. Chem. 2023, 299, 105188, doi:10.1016/j.jbc.2023.105188.

32. Harris, L.A.; Hogg, J.S.; Tapia, J.-J.; Sekar, J.A.P.; Gupta, S.; Korsunsky, I.; Arora, A.; Barua, D.; Sheehan, R.P.; Faeder, J.R. BioNetGen 2.2: Advances in Rule-Based Modeling. Bioinformatics 2016, 32, 3366–3368, doi:10.1093/bioinformatics/btw469.

33. Ritz, C.; Baty, F.; Streibig, J.C.; Gerhard, D. Dose-Response Analysis Using R. PLOS ONE 2015, 10, e0146021, doi:10.1371/journal.pone.0146021.

34. Beal SL; Sheiner LB; Boeckmann AJ; Lb Sheiner *NONMEM Users’ Guides*; ICON Development Solutions: Ellicott City, Maryland, USA, 2016;

35. Claret, L.; Jin, J.Y.; Ferté, C.; Winter, H.; Girish, S.; Stroh, M.; He, P.; Ballinger, M.; Sandler, A.; Joshi, A.;, et al. A Model of Overall Survival Predicts Treatment Outcomes with Atezolizumab versus Chemotherapy in Non–Small Cell Lung Cancer Based on Early Tumor Kinetics. Clin. Cancer Res. 2018, 24, 3292–3298, doi:10.1158/1078-0432.CCR-17-3662.

36. Stein, W.D.; Gulley, J.L.; Schlom, J.; Madan, R.A.; Dahut, W.; Figg, W.D.; Ning, Y.; Arlen, P.M.; Price, D.; Bates, S.E.;, et al. Tumor Regression and Growth Rates Determined in Five Intramural NCI Prostate Cancer Trials: The Growth Rate Constant as an Indicator of Therapeutic Efficacy. Clin. Cancer Res. 2011, 17, 907–917, doi:10.1158/1078-0432.CCR-10-1762.

37. Savic, R.M.; Karlsson, M.O. Importance of Shrinkage in Empirical Bayes Estimates for Diagnostics: Problems and Solutions. AAPS J. 2009, 11, 558–569, doi:10.1208/s12248-009-9133-0.

38. Kim, A.; Cohen, M.S. The Discovery of Vemurafenib for the Treatment of BRAF-Mutated Metastatic Melanoma. Expert Opin. Drug Discov. 2016, 11, 907–916, doi:10.1080/17460441.2016.1201057.

39. Murphy, B.M.; Terrell, E.M.; Chirasani, V.R.; Weiss, T.J.; Lew, R.E.; Holderbaum, A.M.; Dhakal, A.; Posada, V.; Fort, M.; Bodnar, M.S.;, et al. Enhanced BRAF Engagement by NRAS Mutants Capable of Promoting Melanoma Initiation. Nat. Commun. 2022, 13, 3153, doi:10.1038/s41467-022-30881-9.

40. Pratilas, C.A.; Taylor, B.S.; Ye, Q.; Viale, A.; Sander, C.; Solit, D.B.; Rosen, N. ^V600E^ BRAF Is Associated with Disabled Feedback Inhibition of RAF–MEK Signaling and Elevated Transcriptional Output of the Pathway. Proc. Natl. Acad. Sci. 2009, 106, 4519–4524, doi:10.1073/pnas.0900780106.

41. Shin, S.J.; Lee, J.; Kim, T.M.; Kim, J.-S.; Kim, Y.J.; Hong, Y.S.; Kim, S.Y.; Kim, J.E.; Lee, D.H.; Hong, Y.;, et al. A Phase Ib Trial of Belvarafenib in Combination with Cobimetinib in Patients with Advanced Solid Tumors: Interim Results of Dose-Escalation and Patients with NRAS-Mutant Melanoma of Dose-Expansion. J. Clin. Oncol. 2021, 39, 3007–3007, doi:10.1200/JCO.2021.39.15_suppl.3007.

42. Palmer, A.C.; Sorger, P.K. Combination Cancer Therapy Can Confer Benefit via Patient-to-Patient Variability without Drug Additivity or Synergy. Cell 2017, 171, 1678–1691.e13, doi:10.1016/j.cell.2017.11.009.

43. Lehár, J.; Krueger, A.S.; Avery, W.; Heilbut, A.M.; Johansen, L.M.; Price, E.R.; Rickles, R.J.; Short Iii, G.F.; Staunton, J.E.; Jin, X.;, et al. Synergistic Drug Combinations Tend to Improve Therapeutically Relevant Selectivity. Nat. Biotechnol. 2009, 27, 659–666, doi:10.1038/nbt.1549.

