## Supplemental Figure 1 for "Computational modeling of drug response identifies mutant-specific constraints for dosing panRAF and MEK inhibitors in melanoma"

a)

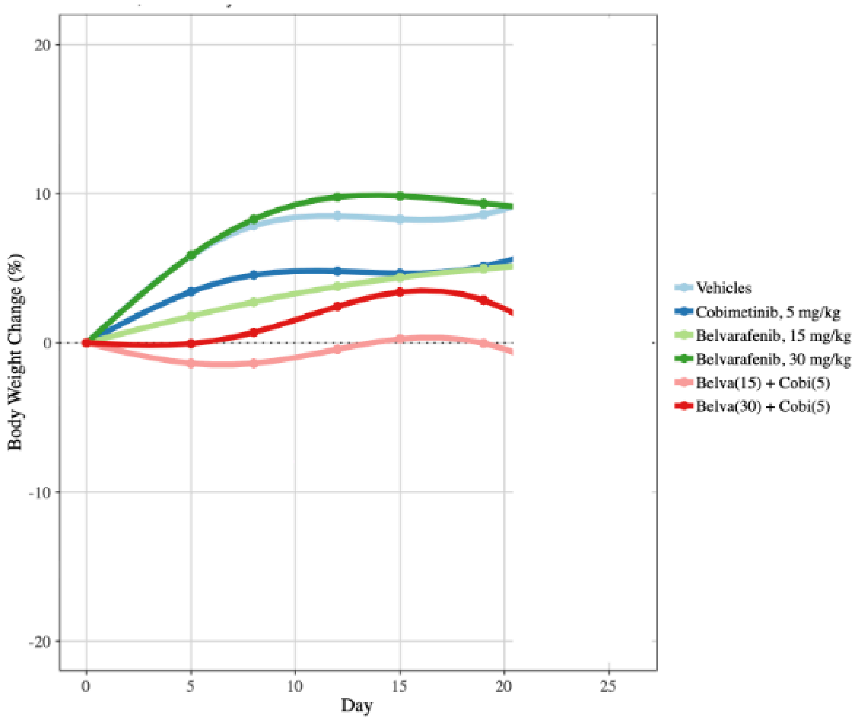


b)

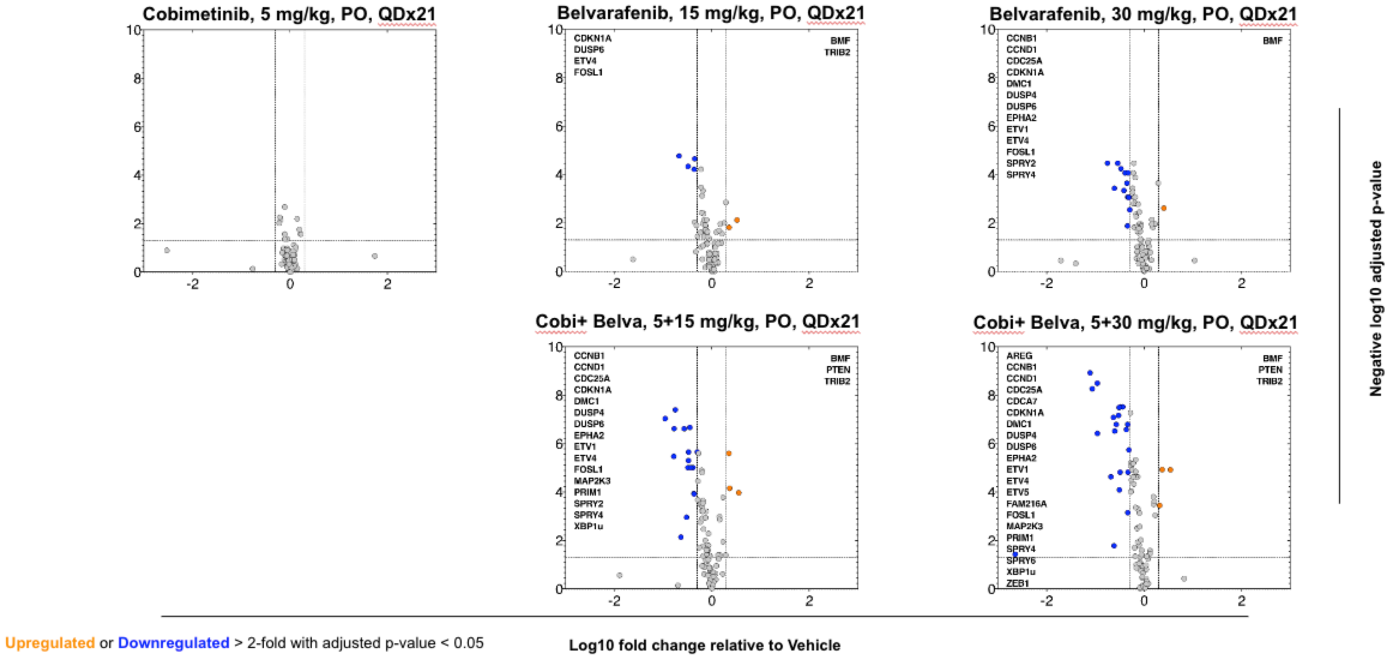


**Figure S1.** a) Body weight of mice in xenograft experiment. b) Gene expression response in IPC-298 xenografts at indicated times and doses of Belvarafenib and Cobimetinib. Upregulated or downregulated genes are marked in color and named on the sides. Three out of five conditions re-analyzed here were previously published in [25].
