## Supplemental Figure 2 for "Computational modeling of drug response identifies mutant-specific constraints for dosing panRAF and MEK inhibitors in melanoma"

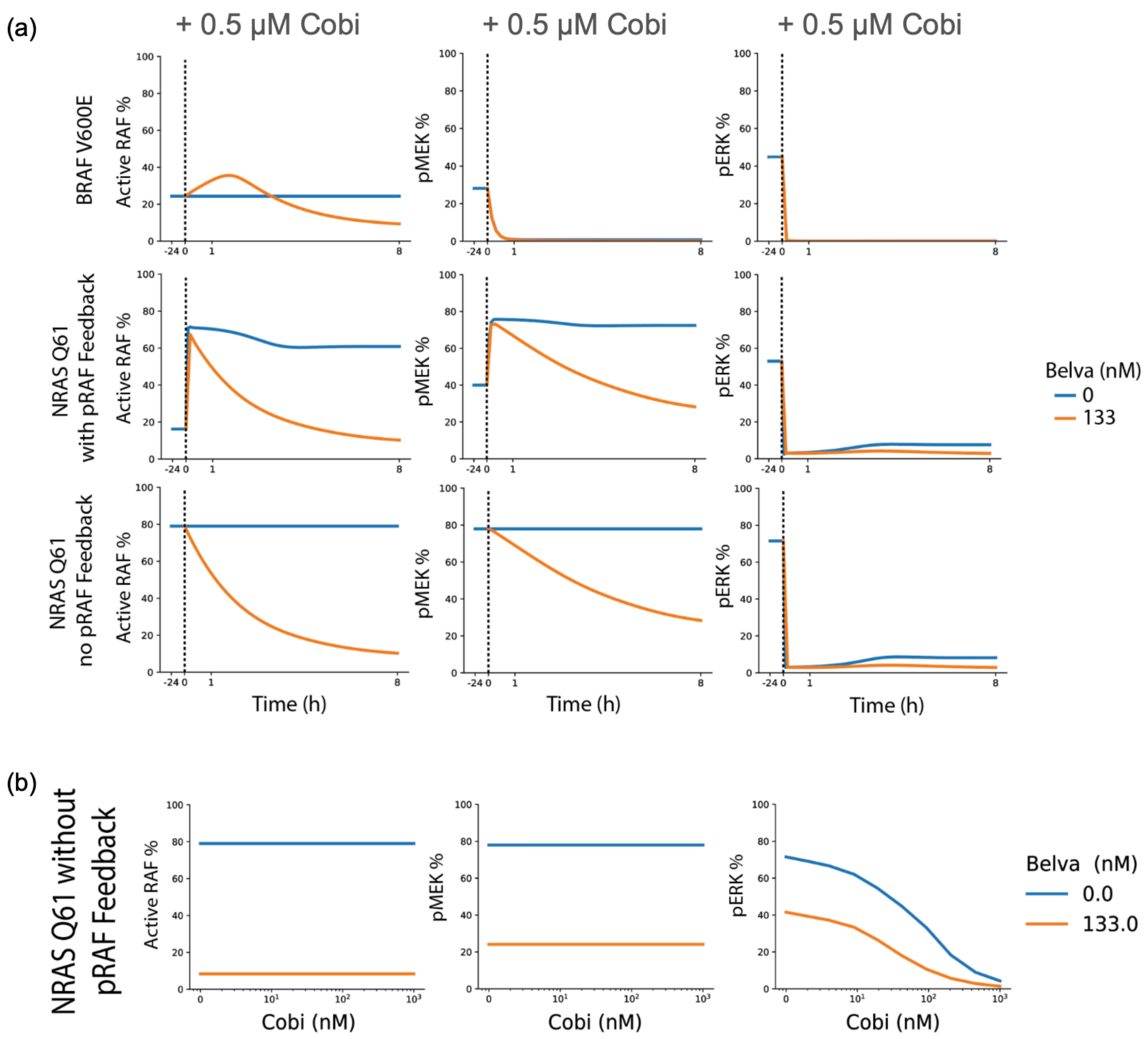


**Figure S2.** Additional model predictions. (a) Model predictions for percentages of active RAF, pMEK, and pERK over time. Models are initially in steady state without drug addition before being dosed with indicated Cobimetinib and Belva concentrations at t = 0. (b) Model predictions for steady state percentages of active RAF, pMEK, and pERK under indicated levels of Belvarafenib and Cobimetinib. Results are shown for NRAS Q61 model without pRAF feedback mechanism.
