## Supplemental Figure 3 for "Computational modeling of drug response identifies mutant-specific constraints for dosing panRAF and MEK inhibitors in melanoma"

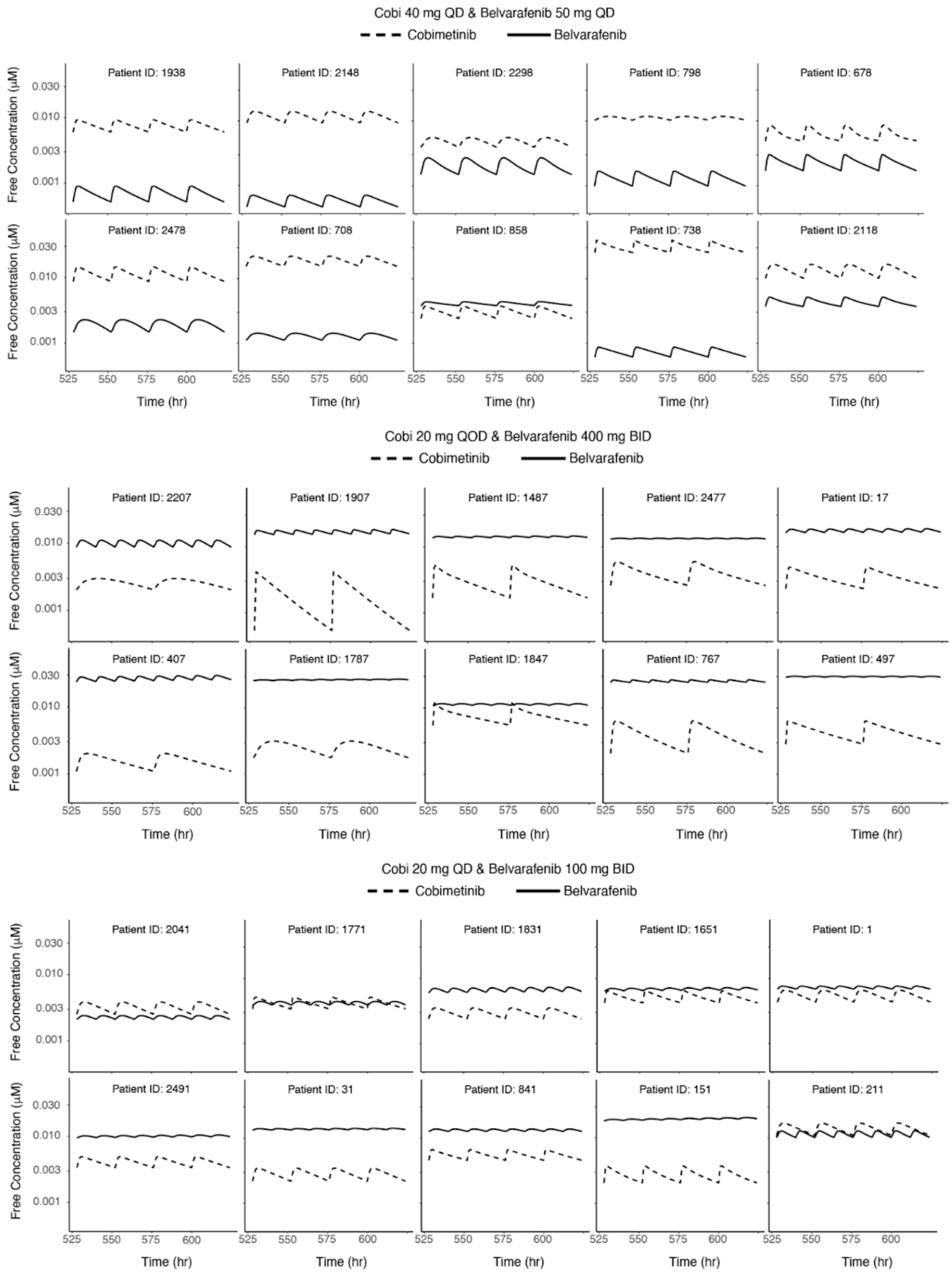


**Figure S3.** PK trajectories corresponding to the drug regimen and virtual patient combinations from fig 6. 96 hours are shown corresponding to at least two complete cycles of drug concentrations, the first 48 hours are used for the analysis shown in fig 6.
