## Supplementary figures and images for "Computational modeling of drug response identifies mutant-specific constraints for dosing panRAF and MEK inhibitors in melanoma"

### Supplemental Figure 4

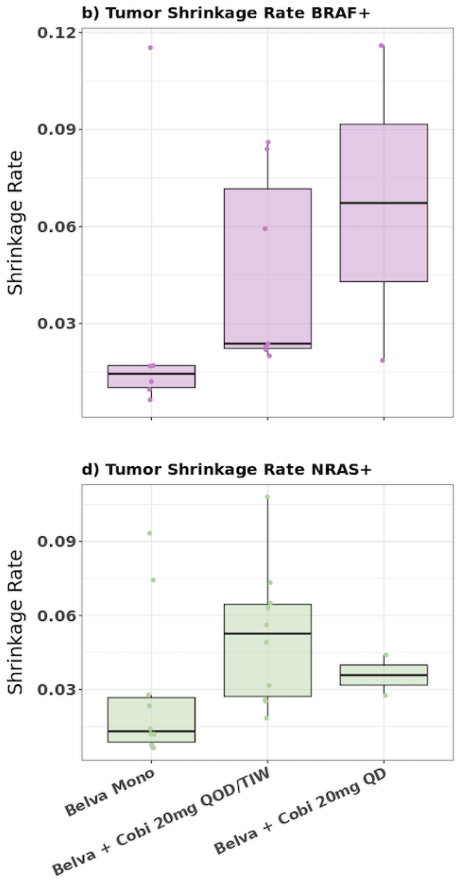


**Figure S4**. Simulations of shrinking rate of tumors in patients.
