## Supplemental Table 1 for "Computational modeling of drug response identifies mutant-specific constraints for dosing panRAF and MEK inhibitors in melanoma"

**SUPPLEMENTAL MATERIAL**

Table S1. Study Design for xenograft experiment.

| *Group* | *No./ Sex* | *Treatment* | *Dose level (mg/kg)[a]* | *No./Sex* | *Route* | *Days of Dosing* | *Dose Conc. (mg/mL)[a]* | *Dose Volume (mL/kg)* |
|---|---|---|---|---|---|---|---|---|
| *1* | *10/F* | *Vehicles* | *0 (Vehicles)* | *10/F* | *PO* | *21* | *0* | *5, 5* |
| *2* | *10/F* | *Cobimetinib* | *5* | *10/F* | *PO* | *21* | *1.1* | *5* |
| *3* | *10/F* | *Belvarafenib* | *15* | *10/F* | *PO* | *21* | *3.3* | *5* |
| *4* | *10/F* | *Belvarafenib* | *30* | *10/F* | *PO* | *21* | *6.6* | *5* |
| *5* | *10/F* | *Belvarafenib + Cobimetinib* | *15 + 5* | *10/F* | *PO* | *21* | *3.3, 1.1* | *5, 5* |
| *6* | *10/F* | *Belvarafenib + Cobimetinib* | *30 + 5* | *10/F* | *PO* | *21* | *6.6, 1.1* | *5, 5* |

*Conc. = concentration; PO = orally; QD = once daily.*

*Note: Vehicle controls were 5% dimethyl sulfide/5% Cremophor EL (100 µ) + 0.5% (w/v) methylcellulose; 0.2% Tween 80™ (100 µL). Dose levels and concentrations are expressed as free-base equivalents and were dosed once daily (QD) for 21 days.*
